# Dissecting the heterogeneity of DENV vaccine-elicited cellular immunity using single-cell RNA sequencing and cellular metabolic profiling

**DOI:** 10.1101/511428

**Authors:** Adam T. Waickman, Kaitlin Victor, Tao Li, Kristin Hatch, Wiriya Rutvisuttinnunt, Carey Medin, Benjamin Gabriel, Richard G. Jarman, Heather Friberg, Jeffrey R. Currier

## Abstract

Generating effective and durable T cell immunity is a critical prerequisite for vaccination against dengue virus (DENV) and other viral diseases. Understanding the precise molecular mechanisms of vaccine-elicited T cell immunity remains a critical knowledge gap in vaccinology. In this study, we utilized single-cell RNA sequencing (scRNAseq) and TCR clonotype analysis to demonstrate that a unique transcriptional signature is present in acutely-activated and clonally-expanded T cells that become committed to the memory repertoire. This effector-associated transcriptional signature is dominated by a unique metabolic transcriptional program. Based on this transcriptional signature, we were able to define a set of functional markers that identify the most potent and durable vaccine-reactive memory precursor CD8^+^ T cells. The transferrin receptor (CD71) other important transporters of amino acids, and direct measurements of glucose and fatty acid uptake, were further validated as early markers of durable T cell memory using conventional flow cytometry. These data suggest that generating durable T cell immunity is a process that is determined by metabolic programming and metabolite availability during the acute phase of the immune response. This study illustrates the power of scRNAseq as an analytical tool to assess the molecular mechanisms of host control and vaccine modality in determining the magnitude, diversity, and persistence of vaccine-elicited cell-mediated immunity.

## INTRODUCTION

An important goal of dengue virus (DENV) vaccine design is to utilize a strategy which induces a broadly effective immune response that encompasses both humoral and cellular immunity. Consisting of four immunologically and genetically distinct serotypes, DENV-1 to DENV-4, DENV infects between 280-500 million individuals yearly worldwide [1–7]. Although the majority of DENV-infected individuals exhibit few or no clinical symptoms, the virus causes approximately 100 million cases of acute febrile illness in humans yearly [1, 6, 7]. While the majority of DENV-infected individuals recover quickly without the need for extensive medical intervention, nearly 500,000 individuals a year develop severe dengue disease, classified as either Dengue Hemorrhagic Fever (DHF) or Dengue Shock Syndrome (DSS) [1, 6, 7]. Characterized by increased vascular permeability, hypovolemia, and dysregulated blood clotting, DHF/DSS has a 2.5% mortality rate [1–7]. The environmental and genetic factors responsible for the development of DHF/DSS are complex and incompletely understood [3, 5–14]. However, prior infection with one serotype of DENV has been shown to significantly increase the likelihood of developing DHF/DSS upon re-infection with a heterotypic viral serotype [5, 9, 15]. This phenomenon is thought to be facilitated at least in part by poorly-neutralizing, serotype cross-reactive antibodies which enable the opsonization of viral particles but not functional neutralization: a process referred to as antibody-dependent enhancement of infection [6, 9, 10, 12, 16]. Hence, the development of an efficacious DENV vaccine has been significantly complicated by the necessity of generating a protective immune response against four distinct serotypes of DENV, while simultaneously avoiding the development of DHF/DSS potentiated via immune-mediated enhancement of infection. A durable and effective cytotoxic T lymphocyte (CTL) response is considered an important component of DENV immunity that may counteract any deleterious effects of cross-reactive antibodies [17–19].

Generating an effective and durable CTL response by vaccination in humans has been largely attributable to live-attenuated vaccines (e.g., vaccinia, yellow fever) and non-replicating viral vectors (rAD/MVA) [20–23]. Despite the wealth of immune monitoring data that has been generated in recent years, a knowledge gap still exists in our understanding of what determines the magnitude, diversity, and persistence of vaccine-elicited CTL responses in humans. Early studies using “systems biology” approaches have focused upon innate immune signatures that correlate with adaptive CD8^+^ T cell responses. Querec et al. found that the induction of expression of the GCN2 kinase by the yellow fever virus (YFV) 17D vaccine at early time points correlated with the frequency of acutely activated (CD38^+^HLA-DR^+^) CD8^+^ T cells at day 15 post-vaccination [24]. However, they did not evaluate the magnitude or function of the memory YFV-specific T cell response induced by the vaccine, nor the associations of their findings with vaccine efficacy. In contrast, similar analysis of the Ad5-vectored HIV vaccine studied in the STEP trial revealed a more rapid up-regulation of innate immune responses than the YFV 17D vaccine, and cytokine responses, especially levels of MCP-2 and MCP-1, correlated with the magnitude of the HIV-specific CD8^+^ T cell response measured by cytokine flow cytometry [25]. More recent studies have focused on studying the relationship between acutely activated CD4^+^ and CD8^+^ T cells after vaccination and during the subsequent convalescent phase. For example, a distinct transcriptional and molecular signature in CD4^+^ T cells was used to distinguish effector phase versus memory responses in BCG vaccinated individuals [26]. Greenough et al. demonstrated that particular transcription factors (eomesodermin and T-bet) allowed the identification of CD8^+^ T cells that expanded rapidly during acute infectious mononucleosis and enabled the proliferation and transition of these cells to later differentiation stages in convalescence [27]. While these studies have revealed distinct transcriptional signatures that correlate early immune activation of antigen-specific T cells with the presence of long-term T cell responses, each has relied upon studying sorted T cells at the population level. An unknown factor is whether antigen-specific T cells that expand rapidly in the effector phase of the immune response within an individual have differential transcriptional profiles that can predict the fate of a particular cell and ultimately its transition into a long-term memory cell. An analytical tool that allows the simultaneous tracking of transcriptional profiles and full-length TCR sequences at the singlecell level would provide an unprecedented understanding of host control in determining the magnitude, diversity, and persistence of vaccine-elicited T cell responses.

TAK-003 is a recombinant, tetravalent DENV vaccine platform derived from the PDK-53 DENV-2 virus strain [28]. Initially derived from the WT DENV-2 16681 isolate, the PDK-53 strain was attenuated by serial passage in primary dog kidney (PDK) cells and has previously been shown to be safe, immunogenic, and capable of stimulating durable cellular and humoral DENV-2 immunity [29–33]. Recombinant viruses were created using the PDK-53 genetic backbone and the precursor membrane (prM) and envelop (E) genes from DENV-1, −3 and −4, resulting in a vaccine product capable of generating an immune response against all four DENV serotypes [34]. This tetravalent formulation was also shown to be safe, immunogenic, and capable of providing protective immunity against DENV challenge in both small animal models and non-human primates [28, 34, 35].

In this study, we demonstrate that TAK-003 elicits a potent cellular immune response which persists for at least 120 days post-vaccination in human subjects. The antigen specificity of the cellular immune response generated by TAK-003 spans the DENV proteome and demonstrates significant crossreactivity against all four DENV serotypes. Single-cell RNA sequencing analysis of CD8^+^ T cells activated in response to TAK-003 exposure revealed a highly polyclonal CD8^+^ T cell repertoire, which had significant clonal overlap between DENV-2 non-structural protein (NS)1- and NS3-reactive CD8^+^ T cells identified and isolated 120 days post-vaccination. Transcriptional analysis of CD8^+^ T cells acutely activated in response to TAK-003 exposure also revealed a highly diverse transcriptional profile, with NS1- and NS3-reactive memory-precursor CD8^+^ T cells at day 14 post-immunization displaying a distinct transcriptional signature dominated by metabolic pathways. Based on these observations, we identified a panel of metabolic markers which could be used to faithfully identify CD8^+^ T cells activated *in vitro* in response to antigenic stimulation, or activated *in vivo* in response to TAK-003 administration. In particular, expression of the transferrin receptor (CD71) – critical for efficient iron uptake - exclusively marks CD8^+^ T cells with high proliferative and effector/memory potential. Therefore, analysis of the metabolic profile in vaccine-responsive CD8^+^ T cells can aid in the identification and characterization of the most effective and durable vaccine-elicited clonotypes.

## RESULTS

### Vaccination with a live-attenuated tetravalent DENV vaccine elicits potent and durable cellular immunity

T cell activation in response to TAK-003 administration was assessed by flow cytometry in 55 individuals on days 0, 14, 28 and 120 post immunization. Consistent with other live-attenuated vaccine platforms, TAK-003 administration resulted in significant CD8^+^ T cell activation on days 14 and 28 post-vaccination (Figure 1A, Figure 1B, **Supplemental Figure 1**). CD8^+^ T cell activation, as assessed by CD38/HLA-DR upregulation, peaked on day 28 post-vaccination and returned to baseline levels by day 120. Moderate CD4^+^ T cell activation was also observed in response to TAK-003 administration (Figure 1C, Figure 1D, **Supplemental Figure 1**), with the peak of activation observed on day 14 post-vaccination.

**Figure 1.**
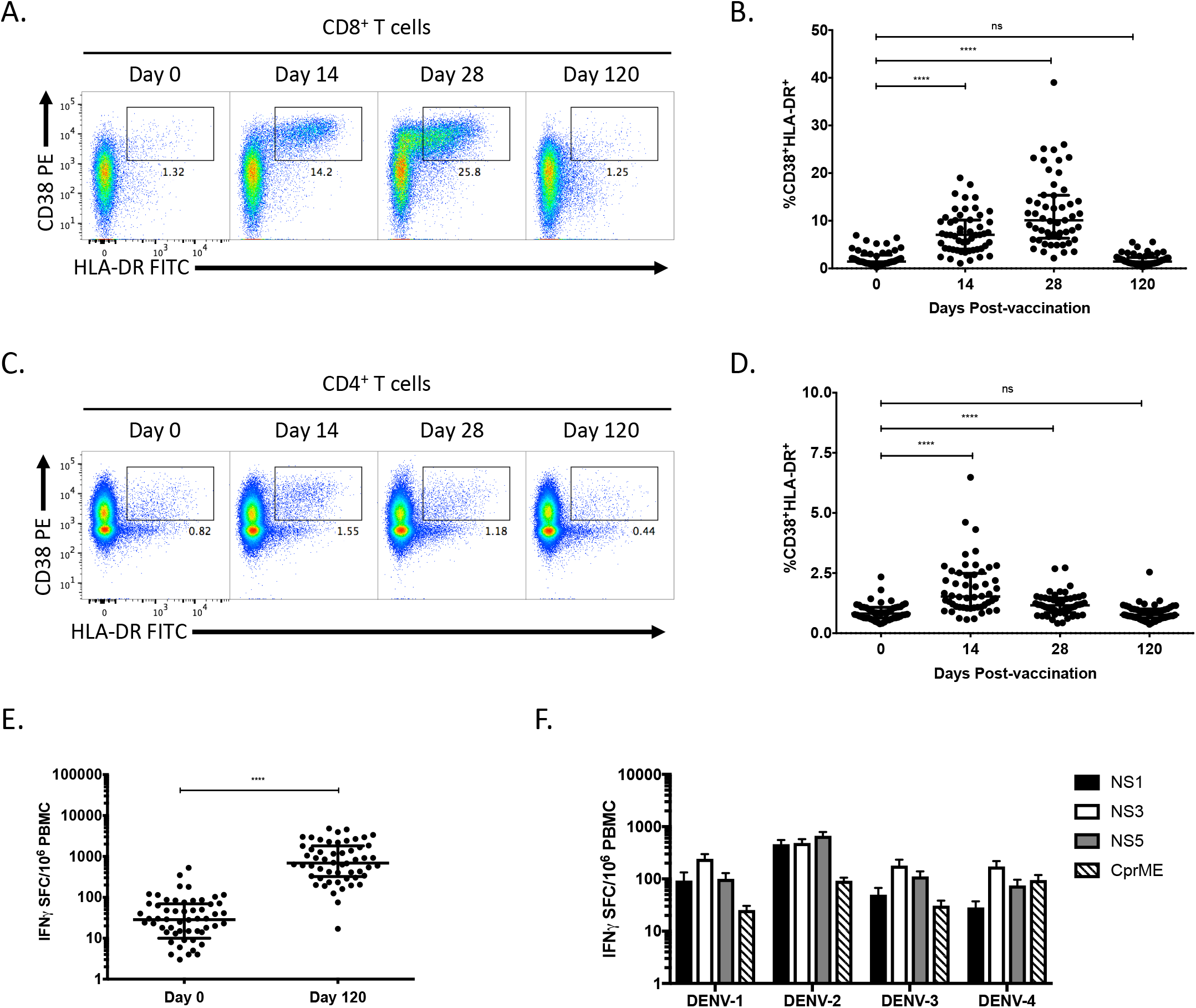
TAK-003 elicits a potent and durable DENV-specific cellular immune response following vaccination. PBMCs from individuals immunized with TAK-003 were assessed for markers of T cell activation by flow cytometry on days 0, 14, 28, and 120 post-vaccination. **A)** Representative plots demonstrating CD8^+^ T cell activation as assessed by CD38 and HLA-DR upregulation on days 0, 14, 28 and 120 post-vaccination. **B)** Aggregate analysis from 55 TAK-003 recipients, demonstrating maximal CD8^+^ T cell activation on day 28 post-vaccination, returning to baseline by day 120. **C)** Representative plots demonstrating CD4^+^ T cell activation as assessed by CD38 and HLA-DR upregulation on days 0, 14, 28 and 120 post-vaccination. **D)** Aggregate analysis from 55 TAK-003 recipients, demonstrating maximal CD4^+^ T cell activation on day 14 post-vaccination, returning to baseline by day 120. **E)** Quantification of total DENV-2-reactive IFN-γ producing T cells from TAK-003-inoculated individuals on days 0 and 120 post-vaccination as determined by ELISPOT. **F)** Antigen-specificity and serotype cross-reactivity of DENV-reactive, IFN-γ producing T cells on day 120 post-vaccination as determined by ELISPOT. N = 55. **** p<0.0001 (Paired t-test)

To determine if the extensive T cell activation observed in response to TAK-003 administration translated to durable cellular immunity, we stimulated PBMCs isolated from study participants on days 0 and 120 of the study with peptide pools corresponding to the NS1, NS3, NS5, and CprM/E proteins of DENV-1 to −4 and quantified the number of vaccine-reactive, IFN-γ producing cells by ELISPOT assay. All subjects receiving TAK-003 displayed a significant increase in the number of circulating vaccine-reactive T cells on day 120 relative to baseline (Figure 1E), with the specificity of this reaction spread across the DENV proteome (Figure 1F). Tetravalent T cell reactivity patterns observed in TAK-003 recipients were detected in both structural (preM/E) and non-structural (NS1-5) regions of the proteome. Structural region responses could have been generated by any, or all, of the four components of the vaccine, however, non-structural responses can be interpreted as truly cross-reactive since DENV-2 is the common non-structural element of the four vaccine components.

### TAK-003 exposure generates a highly polyclonal activated CD8^+^ T cell pool and durable antigen-specific CD8^+^ memory T cells

To further assess the diversity and persistence of TAK-003-elicited CD8^+^ T cell immunity, we utilized single-cell RNA sequencing to track the clonal expansion/contraction of TAK-003-reactive CD8^+^ T cells from acute infection time points and memory time points. To identify and isolate TAK-003-reactive memory CD8^+^ T cells, we stimulated PBMCs from an individual 120 days post TAK-003 administration with DENV-2 derived NS1 and NS3 peptide pools which were identified by ELISPOT analysis as being immunogenic at this time point in this individual (Figure 2A). Following overnight stimulation with the DENV-derived peptide pools, DENV-reactive CD8^+^ T cells were identified by their upregulation of the early activation markers CD25 and CD69. CD8^+^CD25^+^CD69^+^ T cells were isolated by flow cytometric cell sorting (Figure 2B, **Supplemental Figure 2**) and subjected to single-cell RNA sequencing analysis using the 10x Genomics Single Cell Analyzer and human TCR V(D)J reagent system. In addition, vaccine-reactive CD8^+^ T cells from acute-infection time points (day 14) were identified by expression of the activation markers CD38 and HLA-DR and isolated by flow sorting (Figure 2C). A similar number of CD8^+^ CD38^-^HLA-DR^-^ cells from the same sample were isolated for use as a negative control (Figure 2C).

**Figure 2.**
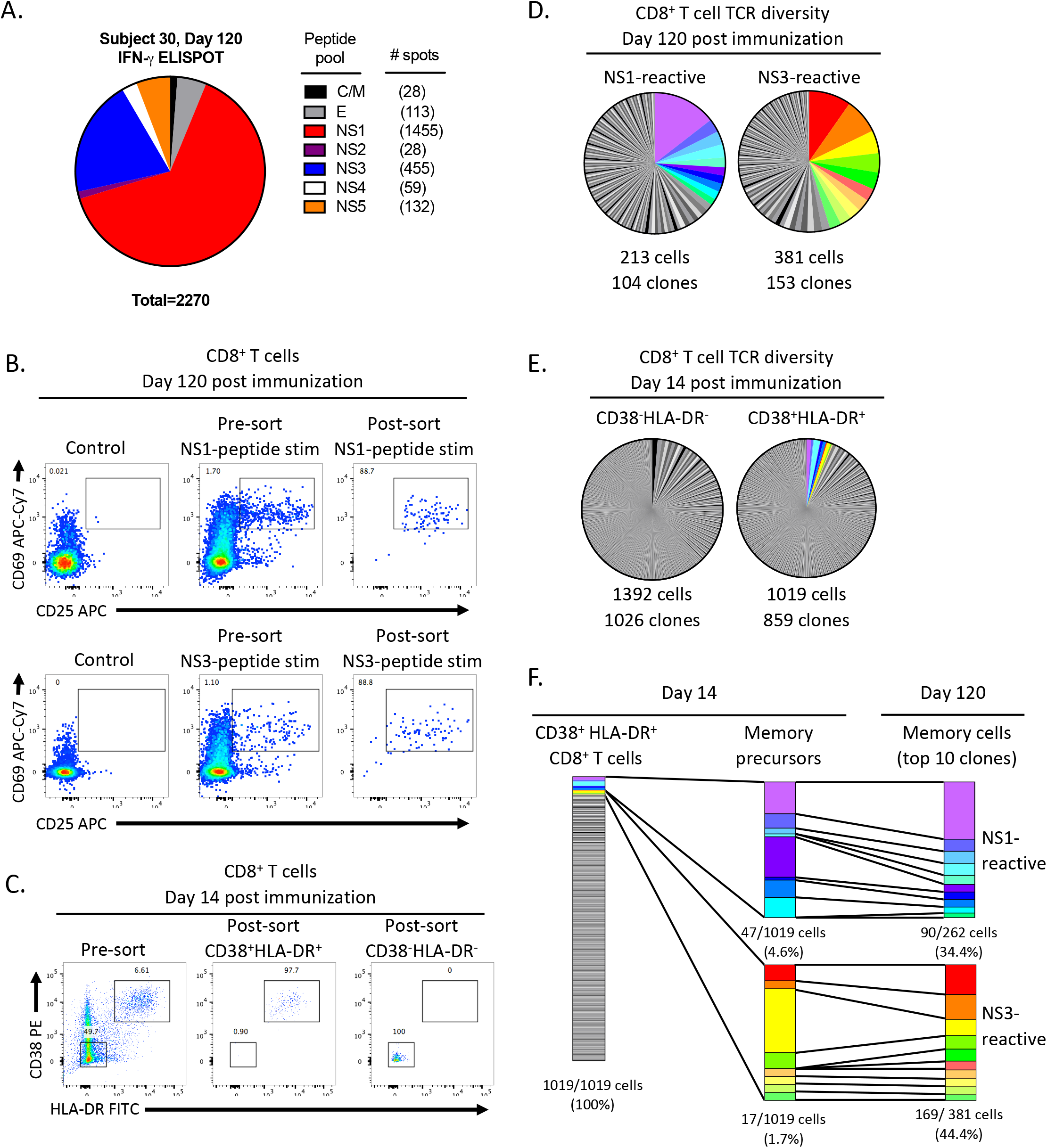
TAK-003-elicited CD38^+^ HLA-DR^+^ CD8^+^ T cells are highly polyclonal and persist as long-lived memory cells. TCR clonotype diversity was assessed within acutely-activated and DENV-reactive memory CD8^+^ T cells using single-cell RNA sequencing. **A)** The antigenic specificity of DENV-reactive memory CD8^+^ T cells from a TAK-003 recipient was assessed by IFN-γ ELISPOT 120 days post immunization. The number of spots is presented relative to 1 million PBMCs. **B)** NS1-reactive and NS3-reactive memory CD8^+^ T cells from 120 post-vaccination were isolated by flow cytometry based on upregulation of CD25 and CD69 expression following *in vitro* stimulation for 18 hours with 1μg/mL of the indicated peptide pools. **C)** Activated (CD38^+^ HLA-DR^+^) CD8^+^ T cells and non-activated (CD38^-^ HLA-DR^-^) CD8^+^ T cells were additionally isolated by flow cytometry from the same TAK-003 recipient 14 day post-vaccination. **D)** TCR clonotype diversity from sorted NS1-reactive and NS3-reactive CD8+ T cells isolated 120 days post-vaccination. The top 10 most abundant NS1- and NS3-reactive clones are demarcated in color. **E)** TCR clonotype diversity in sorted non-activated (CD38^-^ HLA-DR^-^) and activated (CD38^+^ HLA-DR^+^) CD8^+^ T cells using scRNAseq from 14 days post-TAK-003 administration. TCR clones that overlap with the dominant NS1- or NS3-reactive memory CD8+ T cell clones observed at day 120 are indicated with the appropriate color designation. **F)** Assessment of the relative contribution and stability of the dominant NS1- and NS3-reactive memory precursors to the overall CD38^+^HLA-DR^+^ CD8^+^ T cell pool at day 14 post-TAK-003 immunization.

A total of 213 NS1-reactive and 381 NS3-reactive CD8^+^ T cells were identified and isolated from day 120 post-vaccination following *in vitro* stimulation, yielding 104 and 153 unique TCR clonotypes, respectively (Figure 2D, Table 1, **Supplemental Figure 3**). In addition, a total of 1392 CD38^-^HLA-DR^-^ (control) CD8^+^ T cells and 1019 CD38^+^HLA-DR^+^ (vaccine-reactive) CD8^+^ T cells were captured in our single-cell RNA sequencing analysis from day 14 post-vaccination, containing 1026 and 859 unique TCR clonotypes, respectively (Figure 2E, Table 1, **Supplemental Figure 4**). The NS1- and NS3-reactive memory CD8^+^ T cells isolated on day 120 post-vaccination exhibited a significant degree of clonal diversity, with an average (mean) clonal abundance of 1.71 cells/clone for NS1-reactive cells, and 2.49 cells/clone for NS3-reactive cells. However, these statistics are significantly skewed by a large number of clones with a single representative in the dataset. In contrast, the top 10 most abundant clones found in both the NS1- and NS3-reactive memory T cell pool account for 34.4% and 44.4% of recovered cells in the dataset, respectively (Figure 2D, Figure 2F). Furthermore, 80% of these dominant NS1- and NS3-reactive memory CD8^+^ T cell clones identified on day 120 post-vaccination can also be observed in the CD38^+^HLA-DR^+^ CD8^+^ T cell pool isolated on day 14 postvaccination (Figure 2F). Due to practical considerations of sampling it is unlikely that we would capture the entire repertoire of T cells activated acutely following TAK-003 administration. However, even with this limitation, we were still able to capture 80% overlap between the NS1- and NS3-reactive memory clone pool and the CD38^+^HLA-DR^+^ CD8^+^ T cell pool from day 14 post vaccination. The dominant NS1- and NS3-reactive memory precursors account for only 6.3% of all CD38^+^ HLA-DR^+^ CD8^+^ T cells isolated on day 14 post-vaccination (NS1-reactive precursors = 4.6% of CD38^+^HLA-DR^+^ CD8^+^ T cells; NS3-reactive precursors = 1.7% of all CD38^+^HLA-DR^+^ CD8^+^ T cells). Notably, the relative ratio of CD8^+^ T cells at day 14 expressing the dominant NS1- and NS3-reactive TCR clonotypes found at day 120 reflects the relative ratio of NS1 and NS3 responsive cells at day 120 as assessed by ELISPOT. These results suggest that while only a small fraction of the CD38^+^HLA-DR^+^ CD8^+^ T cells elicited in response to vaccination contribute to a persistent pool of memory cells, the relative distribution of antigen reactivity within the vaccine-elicited T cell pool is relativity constant through the course of the response.

**Table 1.**
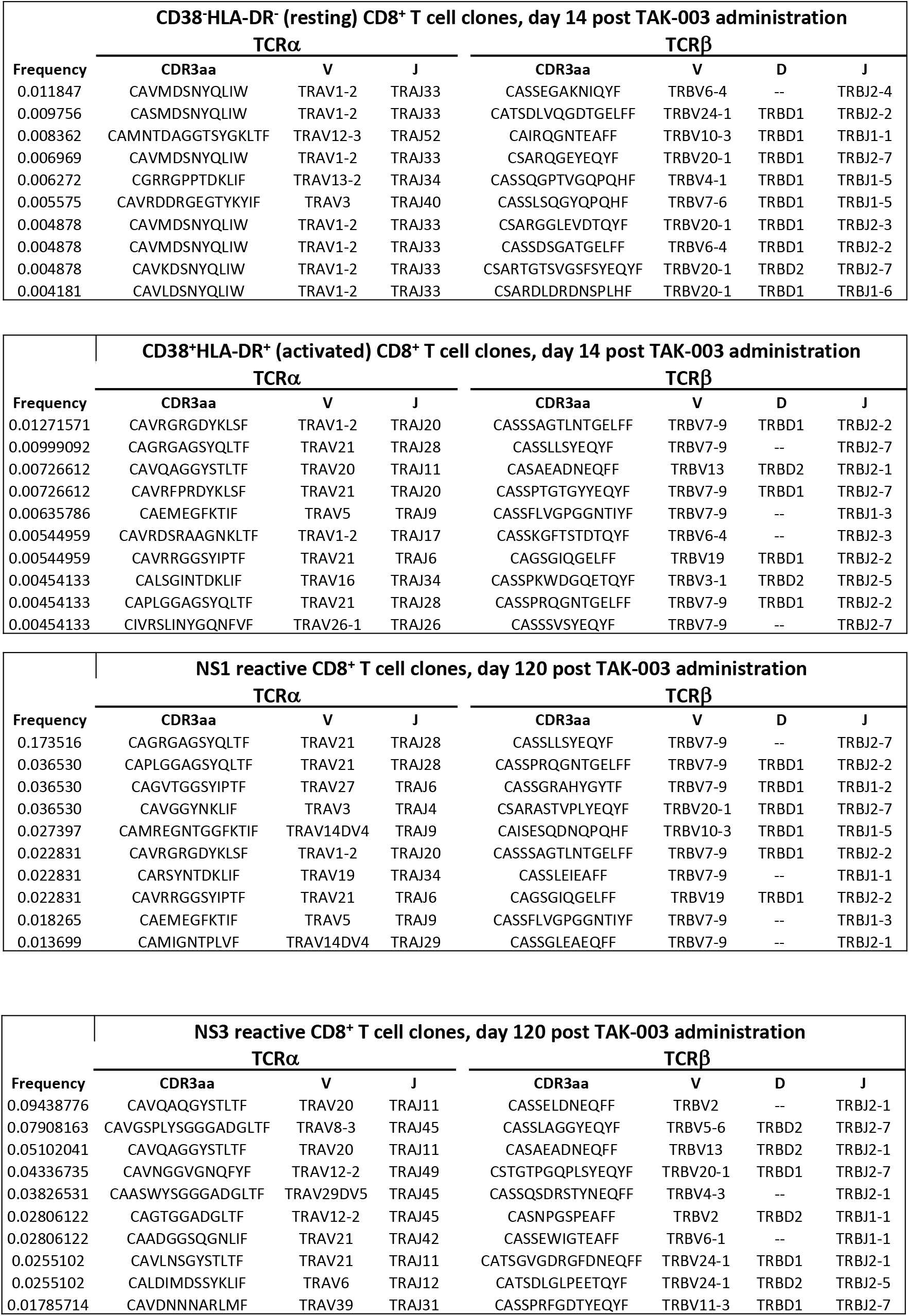
Dominant TCR clonotypes from acutely activated and memory CD8^+^T cells.

### TAK-003-stimulated CD8 T cells exhibit a heterogeneous transcriptional profile, dominated by differential expression of metabolic pathways

In light of the significant clonal diversity observed in the activated CD38^+^HLA-DR^+^CD8^+^ T cell compartment 14 days after TAK-003 administration and the limited number of T cell clones contributing to persistent T cell memory, we decided to determine if the heterogeneity of this population extended to the functional transcriptional profile of these cells. To this end, we assessed the functional gene expression profile of the sorted CD8^+^ CD38^+^HLA-DR^+^ T cells isolated 14 days post-TAK-003 inoculation using single-cell RNAseq. Of the 1019 cells contained within the TCR analysis dataset previously described, we recovered a full gene transcriptional profile from 1003 cells (98.4%) that met the quality control threshold of our analysis pipeline.

Unsupervised tSNE clustering of sorted CD38^+^HLA-DR^+^CD8^+^ T cells based on differential gene expression profiles revealed four statistically-distinct populations (Figure 3A, Table 2). Expression of genes associated with an effector CD8^+^ T cell program such as granzyme A, granzyme B, and perforin were significantly enriched in cluster 1 (Figure 3B, Table 2). Transcripts associated with a more resting/naïve T cell phenotype such as CCR7, MAL, and TCF7 were significantly enriched in clusters 2 and 3, and showed minimal overlap with cells expressing effector gene products (Figure 3B, Table 2). In addition, gene expression associated with cellular metabolism, proliferation and cell-cycle progression such as TYMS, IDH2, and GAPDH were significantly enriched in cluster 1 relative to all other groups.

**Figure 3.**
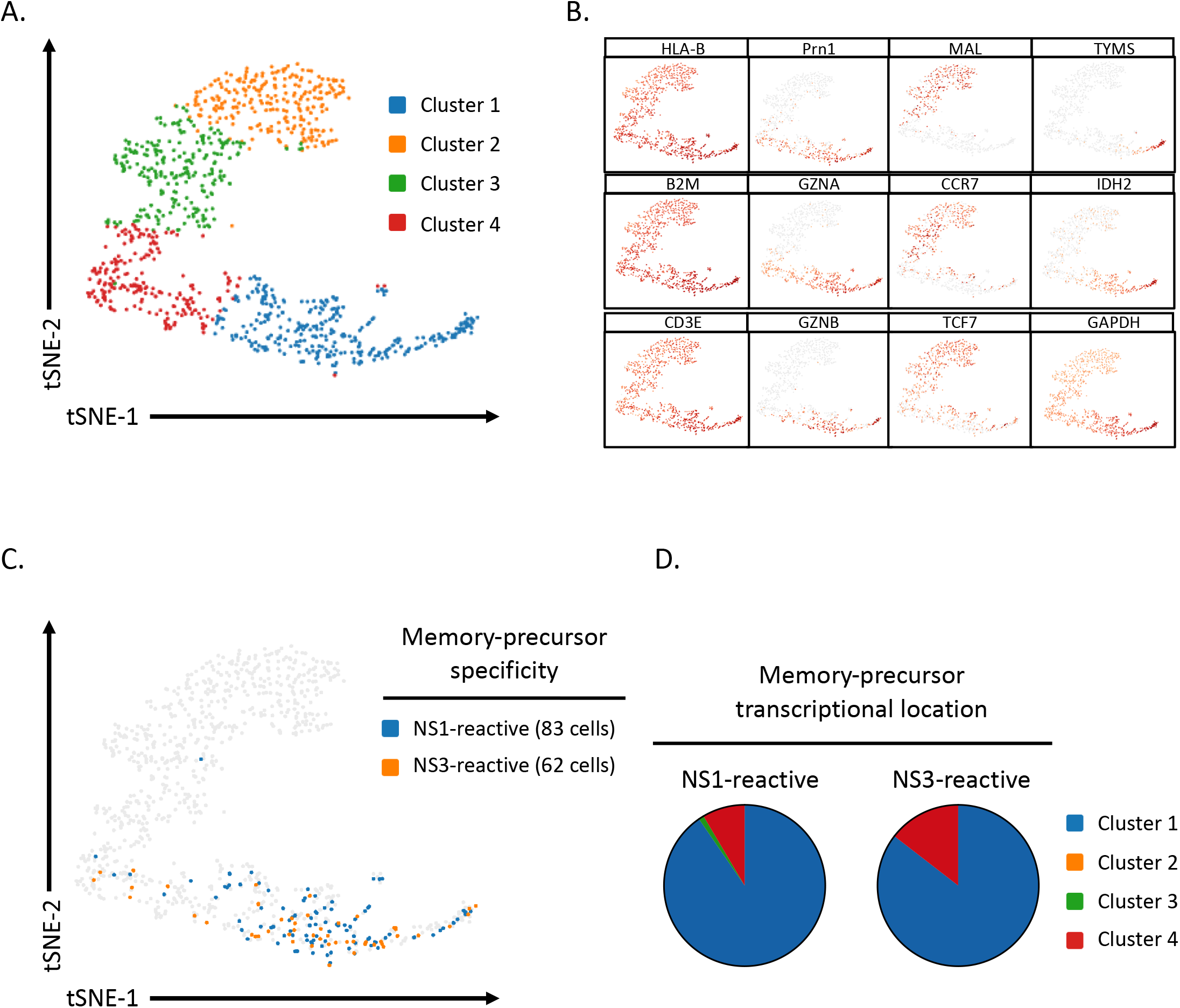
TAK-003-elicited CD8^+^ memory precursors exhibit a unique transcriptional profile. **A)** Transcriptional heterogeneity of sorted CD38^+^HLA-DR^+^ CD8^+^ T cells isolated 14 days post-TAK-003 administration as assessed by single-cell RNA sequencing. Unsupervised cell clustering and data visualization were performed using sparse nearest-neighbor graphing, followed by Louvain Modularity Optimization. **B)** Expression of select transcripts within the sorted CD38^+^HLA-DR^+^ CD8^+^ T cells. **C)** Distribution of CD38^+^HLA-DR^+^ CD8^+^ T cells at day 14 with TCR clonotypes overlapping with NS1- and NS3-reactive memory CD8^+^ T cells at day 120 (memory precursors). **D)** Transcriptional cluster localization of NS1- and NS3-reactive memory precursors at day 14 post-TAK-003 administration.

**Table 2.**
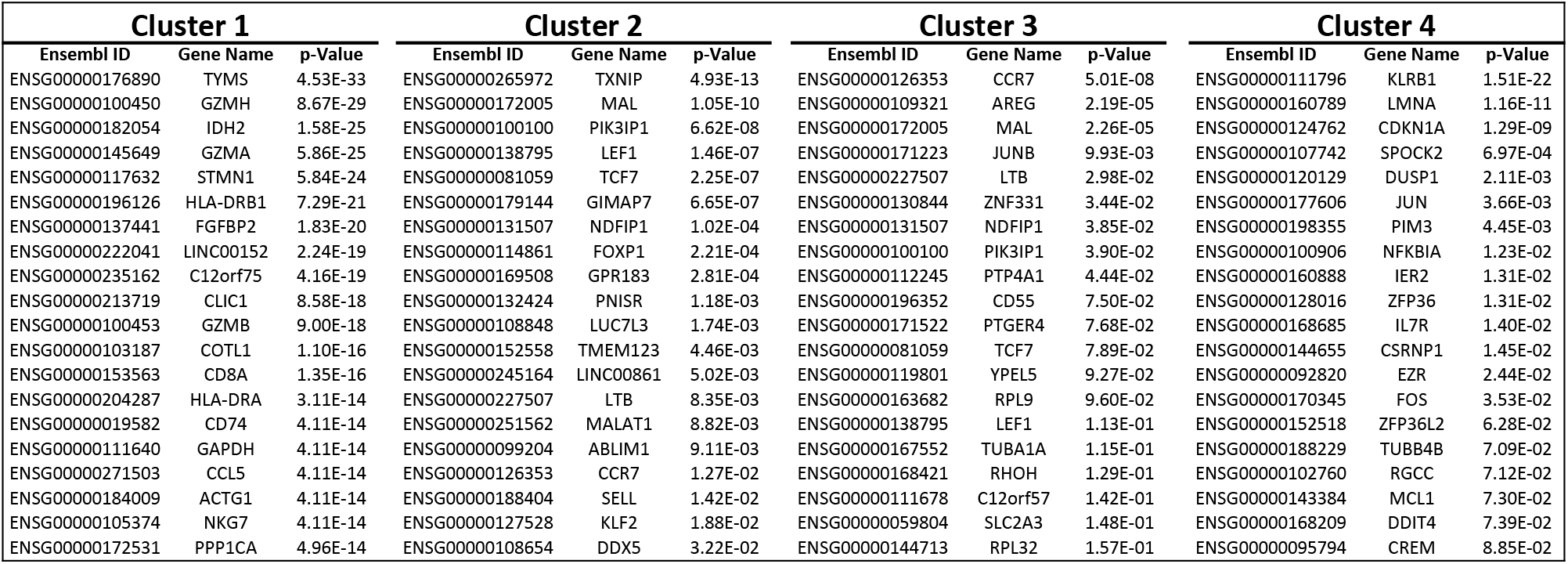
Cluster defining gene lists from CD8^+^ T cell scRNAseq analysis.

Of the 1003 CD38^+^HLA-DR^+^ CD8^+^ T cells recovered on day 14 post-vaccination, 145 cells (14.5%) expressed TCRs found in NS1- and NS3-reactive memory cells on day 120 post-vaccination (Figure 3C). The transcriptional profile of these memory precursors positioned them predominantly within the previously defined phenotypic cluster 1 (Figure 3D), suggesting that this distinct subset of CD38^+^HLA-DR^+^ CD8^+^ T cells found 14 days post-vaccination is uniquely primed to develop into long-lived memory cells.

To further define the transcriptional and phenotypic signature of vaccine-reactive CD8^+^ T cells within our dataset, we utilized the Ingenuity Pathway Analysis (IPA) software package [36] to identify gene pathways selectively expressed in putative DENV-reactive effector/memory-precursor CD8^+^ T cells (Table 3). To this end, we assessed the gene pathways preferentially expressed within cluster 1, which contained majority of identified memory-precursor cells, relative to all other cells in the dataset. Interestingly, there was some preferential expression of gene pathways associated with effector function and cellular migration in cluster 1. However, the dominant cellular transcriptional signatures that distinguished cluster 1 from the rest of the dataset were associated with cellular metabolism and proliferation, such as oxidative phosphorylation, mTOR signaling, and eIF4/p70S6K signaling (Table 3). These data suggest that the assessment of cellular metabolism pathways may provide a robust and unbiased indication of cellular memory-precursor potential, as well as effector status and antigen reactivity.

**Table 3.** Enriched gene pathways in memory-precursor CD8^+^T cells.

| **Pathway** | **p-value** | **z score** | **Ratio** |
| --- | --- | --- | --- |
| Oxidative Phosphorylation | 1.00E-36 | 4.667 | 0.346 |
| EIF2 Signaling | 1.00E-32 | -4.146 | 0.202 |
| Mitochondrial Dysfunction | 3.16E-31 | - | 0.23 |
| mTOR Signaling | 2.51E-16 | 0.816 | 0.137 |
| Regulation of eIF4 and p70S6K signaling | 6.31E-15 | - | 0.149 |
| Sirtuin Signaling Pathway | 6.31E-15 | 0.654 | 0.106 |
| Regulation of Actin-based motility by Rho | 2.00E-13 | 4 | 0.198 |
| Signaling by Rho Family GTPases | 5.01E-12 | 2.836 | 0.099 |
| RhoGDI Signaling | 7.94E-12 | -3.299 | 0.119 |
| Cdc42 Signaling | 2.00E-11 | 2.13 | 0.14 |

### Polyclonal *in vitro* T cell activation leads to an increase in metabolite transporter expression and substrate utilization

To explore the broader implications of the observed relationship between metabolic gene activity and effector/memory potential in vaccine-reactive T cells, we aimed to establish a set of markers to quantify the metabolic potential of human T cells following TCR engagement. With this in mind, we stimulated PBMCs from normal healthy donors *in vitro* with αCD3/CD28 to induce polyclonal T cell activation and assessed the expression of CD71 (transferrin receptor) and CD98 (large neutral-amino acid transporter)-both transmembrane metabolite transporters previously demonstrated to play a critical role in maintaining lymphocyte metabolic homeostasis and effector capability [37–40]. In addition, we directly assessed the ability of *in vitro* activated T cells to utilize glucose (assessed by uptake of 2-NBDG, a fluorescent analog of glucose) or fatty acids (assessed by BODIPY FL-C_16_ uptake, a florescent palmitate derivative) by flow cytometry.

After *in vitro* stimulation, CD4^+^ and CD8^+^ T cells showed a well-characterized pattern of activation marker upregulation (Figure 4A, Figure 4B, **Supplemental Figure 5**). Furthermore, we observed a dramatic and sustained increase in expression of CD71 and CD98 following TCR stimulation (Figure 4C, Figure 4D, **Supplemental Figure 5**), as well as 2-NBDG and BODIPY FL-C_16_ uptake (Figure 4E, Figure 4F, **Supplemental Figure 5**). Of particular note, the expression of CD71 and CD98 and uptake of BODIPY FL-C_16_ showed exceptional promise as markers of T cell activation and metabolic potential due to their large dynamic range and persistence relative to other markers of T cell activation. In particular, the expression of CD71 on *in vitro* stimulated CD8^+^ T cells increased ~600 fold 48 hours after polyclonal stimulation.

**Figure 4.**
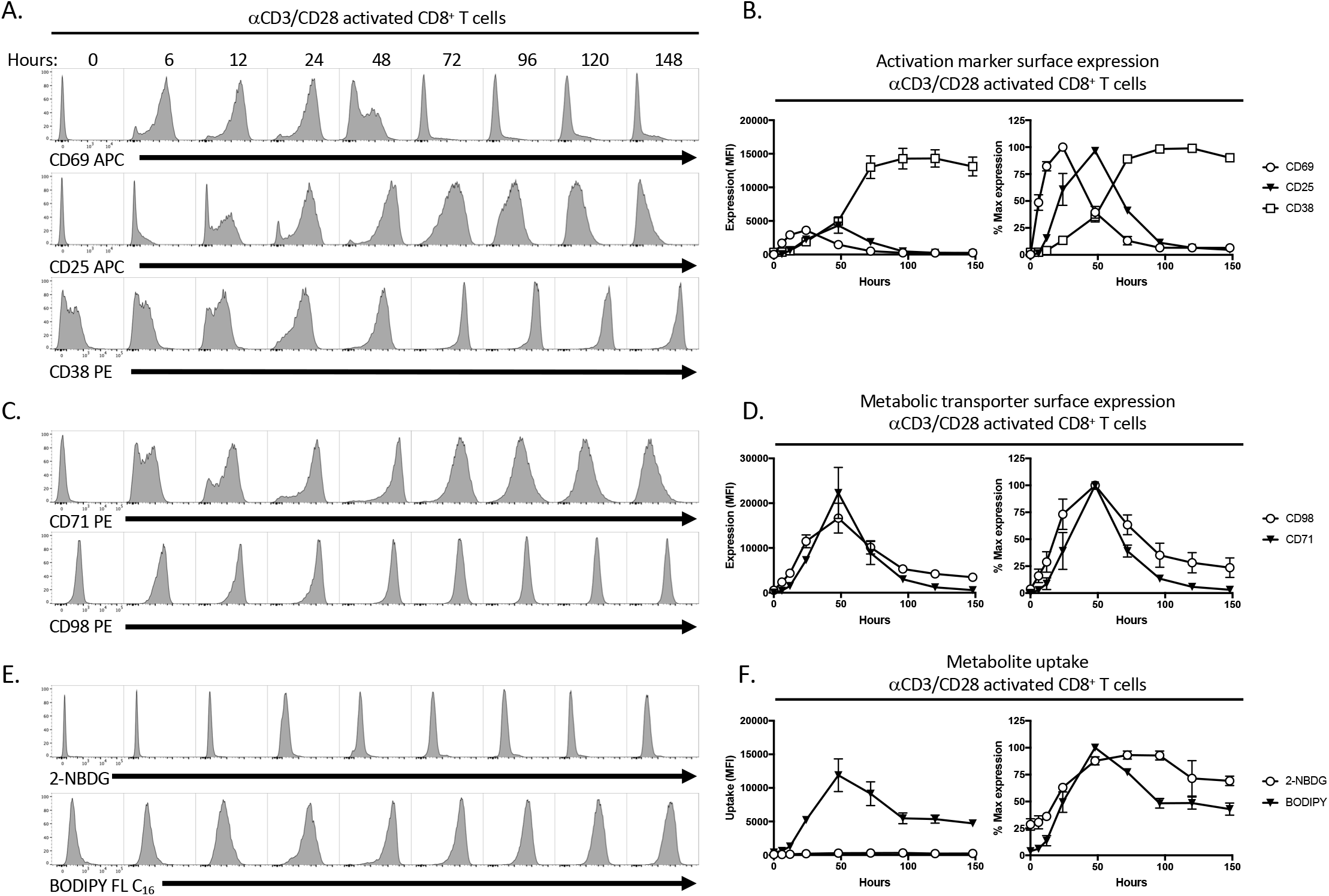
*In vitro* activated T cells can be identified by changes in metabolite transporter expression and metabolite utilization. CD8^+^ T cells from healthy donors were analyzed by flow cytometry at the indicated time points after *in vitro* stimulation with 0.1μg/mL αCD3 and 1μg/mL αCD28. T cell activation was assessed based on expression of **A), B)** CD69 CD25, **C), D)** CD71 CD98, and **E), F)** 2-NBDG BODIPY FL-C16 uptake. Error bars show mean and SEM. Results are representative of two independent experiments with a total of 4 individual donors.

### Antigen-specific *in vitro* T cell activation results in changes in metabolite transporter expression and substrate utilization

In light of the observation that the expression of CD71 and CD98 and the uptake of BODIPY FL-C_16_ are quantifiable and robust metabolic indicators of polyclonal T cell activation following *in vitro* activation, we endeavored to determine if these markers could be utilized to identify and characterize T cells activated in an antigen-specific fashion. For this reason, we stimulated PBMCs from healthy donors with peptide pools corresponding to the proteomes of common viral pathogens (hCMV, HBV, adenovirus, and influenza), and assessed the upregulation of conventional activation markers (CD69, CD25), metabolite transporters (CD71, CD98), as well as the utilization of metabolic substrates (BODIPY FL-C_16_) by flow cytometry.

As anticipated, we observed significant donor-to-donor variability in the fraction of CD8^+^ T cells responding to virus-antigen stimulation by upregulating CD25/CD69 expression (Figure 5A, Figure 5B). Consistent with the demographics of the PBMC donor pool utilized in the experiment, we observed the most abundant CD8^+^ T cell activation in response to CMV peptide stimulation, moderate activation in response to adenovirus and influenza, while minimal activation was seen in response to HBV peptide stimulation. In addition to upregulating expression of CD69 and CD25 following *in vitro* antigenic stimulation, we observed that CD8^+^ T cells concurrently increased their uptake of BODIPY FL-C_16_ (Figure 5A, Figure 5C), expression of CD98 (Figure 5A, Figure 5D), as well as expression of CD71 (Figure 5A, Figure 5E). There was a significant degree of correlation between the upregulation of the conventional activation makers CD25/CD69 following *in vitro* stimulation and the increased expression of CD71, CD98, or the uptake of BODIPY FL-C_16_ following hCMV (Figure 5F), adenovirus (Figure 5G), or influenza (Figure 5H) peptide stimulation in these same cells. These data demonstrate that the antigen-restrictive activation of clonal CD8^+^ T cell can be quantified and functionally characterized using a combination of conventional activation markers, metabolic transporters, and fluorescent metabolic substrates.

**Figure 5.**
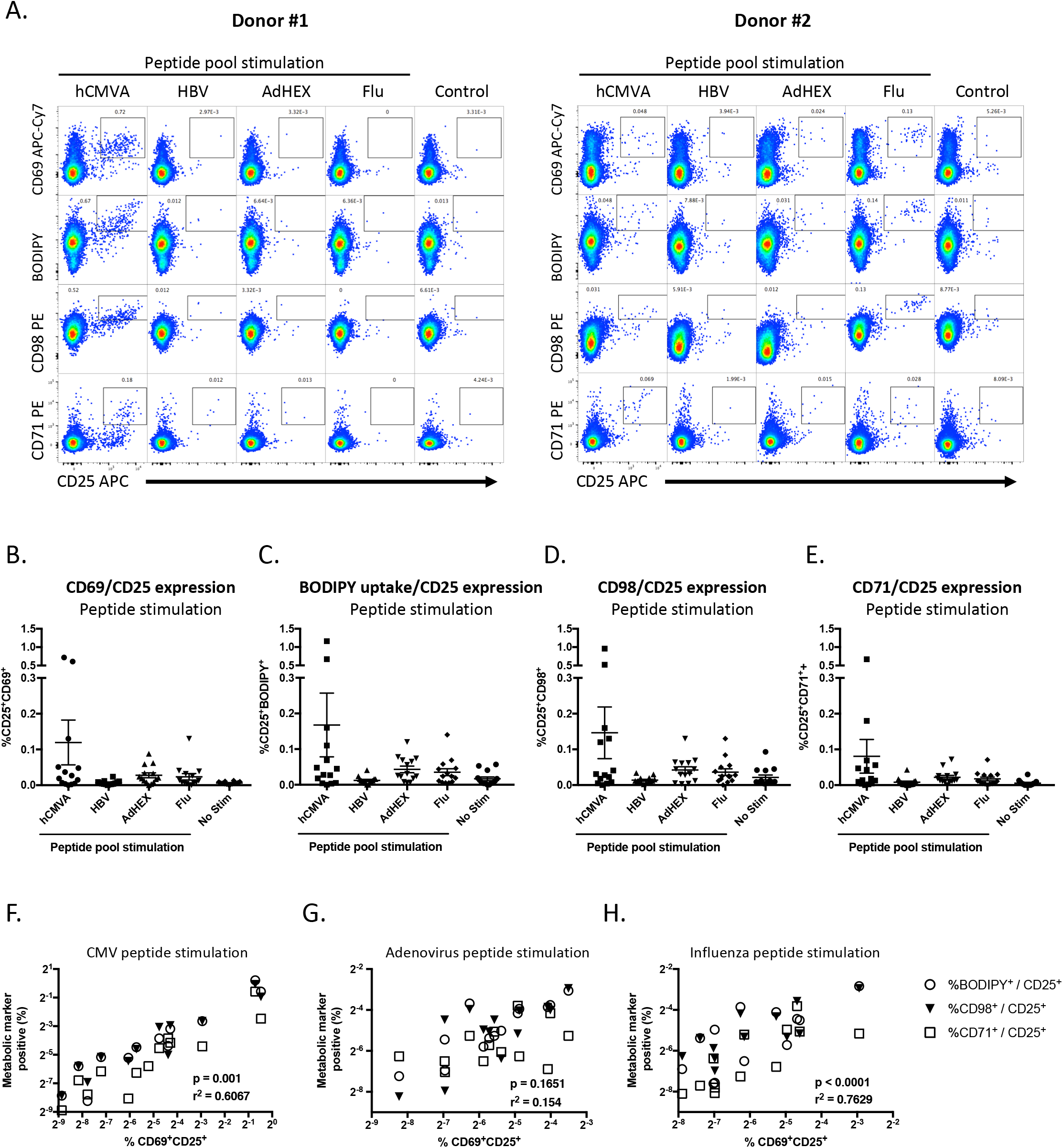
Antigen-specific *in vitro* CD8^+^ T cell activation results in increased metabolite transporter expression and substrate utilization. **A)** CD8^+^ T cells from healthy donors were analyzed by flow cytometry after 48 hours *in vitro* stimulation with 1μg/mL of the indicated peptide pool. T cell activation was assessed based on **B)** upregulation of CD69 and CD25 expression, **C)** increased uptake of BODIPY FL-C_16_ and increased CD25 expression, **D)** increased expression of CD98 and CD25, and **E)** increased expression of CD71 and CD25. Error bars show mean and SEM. The percentage of CD8 T cells exhibiting a metabolically activated state as assessed by BODIPY FL-C16 uptake, CD98 expression, and CD71 expression following *in vitro* stimulation with **F)** hCMV **G)** adenovirus, or **H)** influenza derived peptide stimulation was highly correlated with the percentage of cells expressing the conventional activation markers CD69 and CD25 at the same time point. Results are representative of two independent experiments with a total of 15 individual donors.

### *In vivo* activated T cells can be identified by changes in metabolite transporter expression and metabolite utilization

Having established the utility of CD71/CD98 expression, as well as BODIPY FL-C_16_ and 2-NBDG uptake, for the identification and functional characterization of antigen-specific *in vitro* activated T cells, we aimed to determine whether these same markers can be used to dissect the functional heterogeneity of *in vivo* activated, vaccine-reactive T cells. To this end, we evaluated the expression of the conventional activation markers CD38/HLA-DR on T cells isolated from individuals inoculated with TAK-003 on days 0, 14, 28 and 120 post-immunization, the ability of cells to uptake 2-NBDG and BODIPY FL-C_16_, and the upregulation of the metabolic transporters CD71 and CD98.

As previously shown, TAK-003 administration elicits a potent CD8^+^ T cell response as assessed by CD38/HLA-DR expression, with the peak of CD8^+^ T cell expansion observed on day 28 postinoculation (Figure 6A, **Supplemental Figure 6**). A similar, albeit reduced, pattern of CD4^+^ T cell activation can be concurrently observed in vaccinated individuals, with the peak of CD4^+^ T cell expansion occurring 14 days post-inoculation (**Supplemental Figure 7**).

**Figure 6.**
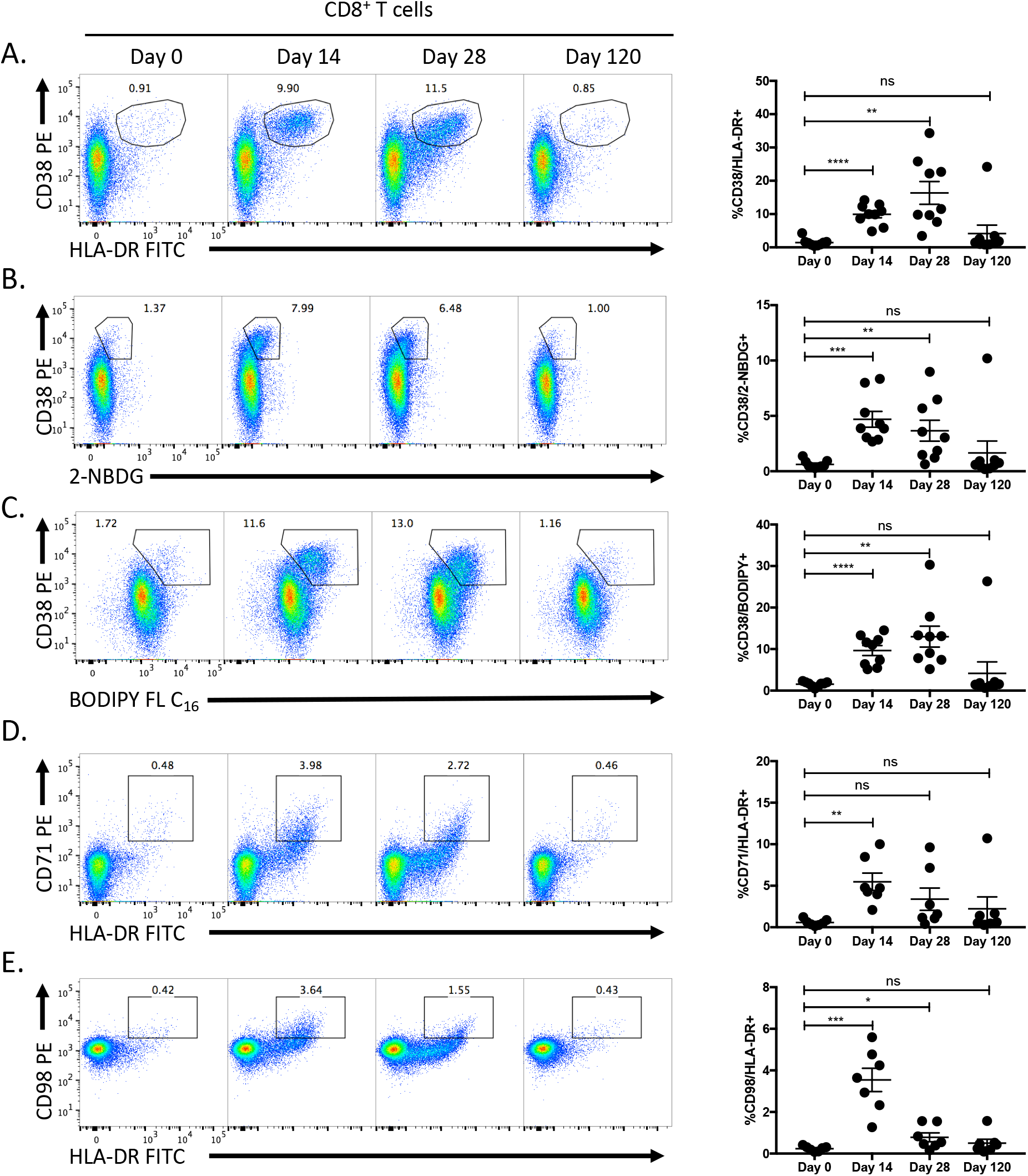
Vaccine-reactive T cells can be identified by changes in metabolite transporter expression and metabolite utilization. CD8^+^ T cells from TAK-003 recipients were analyzed by flow cytometry at days 0, 14, 28 and 120 post-vaccination. Vaccine-reactive CD8^+^ T cells were quantified based on expression of **A)** CD38/HLA-DR, **B)** CD38/2-NBDG, **C)** CD38/BODIPY FL-C_16_, **D)** CD71/HLA-DR, and **E)** CD98/HLA-DR. Error bars show mean and SEM. N = 10 individuals. * p <0.05, *** p<0.001, **** p<0.0001 (Paired t-test)

Consistent with the single-cell RNA sequencing phenotype observed in TAK-003-reactive T cells and our *in vitro* T cell stimulation data, a significant increase in both 2-NBDG and BODIPY FL-C_16_ uptake can be observed in HLA-DR positive CD8^+^ (Figure 6B, Figure 6C) and CD4^+^ (**Supplemental Figure 7**) T cells on days 14 and 28 post-vaccination. Additionally, a distinct subset of HLA-DR positive CD8^+^ and CD4^+^ T cells upregulate CD71 (Figure 6D, **Supplemental Figure 7**) and CD98 (Figure 6E, **Supplemental Figure 7**) on days 14 and 28 post-TAK-003 administration. Notably, while the peak expression of CD38/HLA-DR on CD8^+^ T cells occurs on day 28 post-vaccination, the expression of both CD71 and CD98 peaks on day 14, suggesting that these markers may provide an earlier indicator of vaccine T cell immunogenicity than conventional surface markers. The expression of the conventional activation markers CD38 and HLA-DR and the uptake of BODIPY FL-C_16_ and 2-NBDG - appear to occur concomitantly and uniformly in *in vivo* activated T cells whereas the expression of CD71 (transferrin receptor) shows a significant amount of variability within the HLA-DR^+^ T cell compartment (Figure 6E). As quantification of CD38, CD71, and CD98 was all performed using antibodies conjugated to same fluorophore (PE), the observed heterogeneity in CD71 expression within the HLA-DR^+^ T cell compartment cannot be attributed to a technical aberration. In other words, while the ability to uptake glucose and fatty acids – as assessed by 2-NBDG and BODIPY-FL C16 uptake, respectively - are potential universal indicators of T cell activation, iron metabolism - as assessed by CD71 expression – might more accurately demark those T cells with the highest metabolic demands. This suggests the possibility that heterogeneity of CD71 expression may reflect and capture the functional diversity previously observed in the scRNAseq-derived transcriptional profiles of vaccine-reactive CD8^+^ T cells, and can be utilized to identify effector/memory-precursor CD8^+^ T cells.

### Vaccine-elicited variation in transferrin receptor (CD71) expression correlates with CD8^+^ T cell effector/memory potential

To investigate whether differences in the surface expression of CD71 on *in vivo* activated CD8^+^ T cells can be used to determine their effector/memory potential, we utilized scRNAseq to assess the relative abundance of memory precursor CD8^+^ T cell clonotypes within either the CD71^+^HLA-DR^+^ or CD38^+^HLA-DR^+^ CD8^+^ T cell compartment 14 days post-TAK-003 administration. To this end, CD71^+^HLA-DR^+^ CD8^+^ T cells from the same individual used in the previous scRNAseq analysis were sorted 14 days post-TAK-003 inoculation and subjected to scRNAseq analysis (Figure 7A). This analysis resulted in the capture of 117 individual TAK-003-reactive cells with 80 unique TCR clonotypes (Figure 7B, **Supplemental Table 1**). NS1- and NS3-reactive memory precursors within the sorted CD71^+^HLA-DR^+^ CD8^+^ T cell pool were identified by the presence of TCR clonotypes found in NS1- and NS3-stimulated memory CD8^+^ T cells 120 days post-vaccination (Figure 2B, Figure 2D, Table 1). Of the 117 captured CD71^+^HLA-DR^+^ CD8^+^ cells within the scRNAseq dataset, 31 and 24 were NS1- or NS3-reactive memory precursors, respectively (Figure 7B). These numbers represent 47% of all cells within the sorted population: ~4 fold higher than the frequency of the same TCR clonotypes within the sorted CD38^+^HLA-DR^+^ CD8^+^ T cell pool previously analyzed (Figure 2D, Figure 7B).

**Figure 7.**
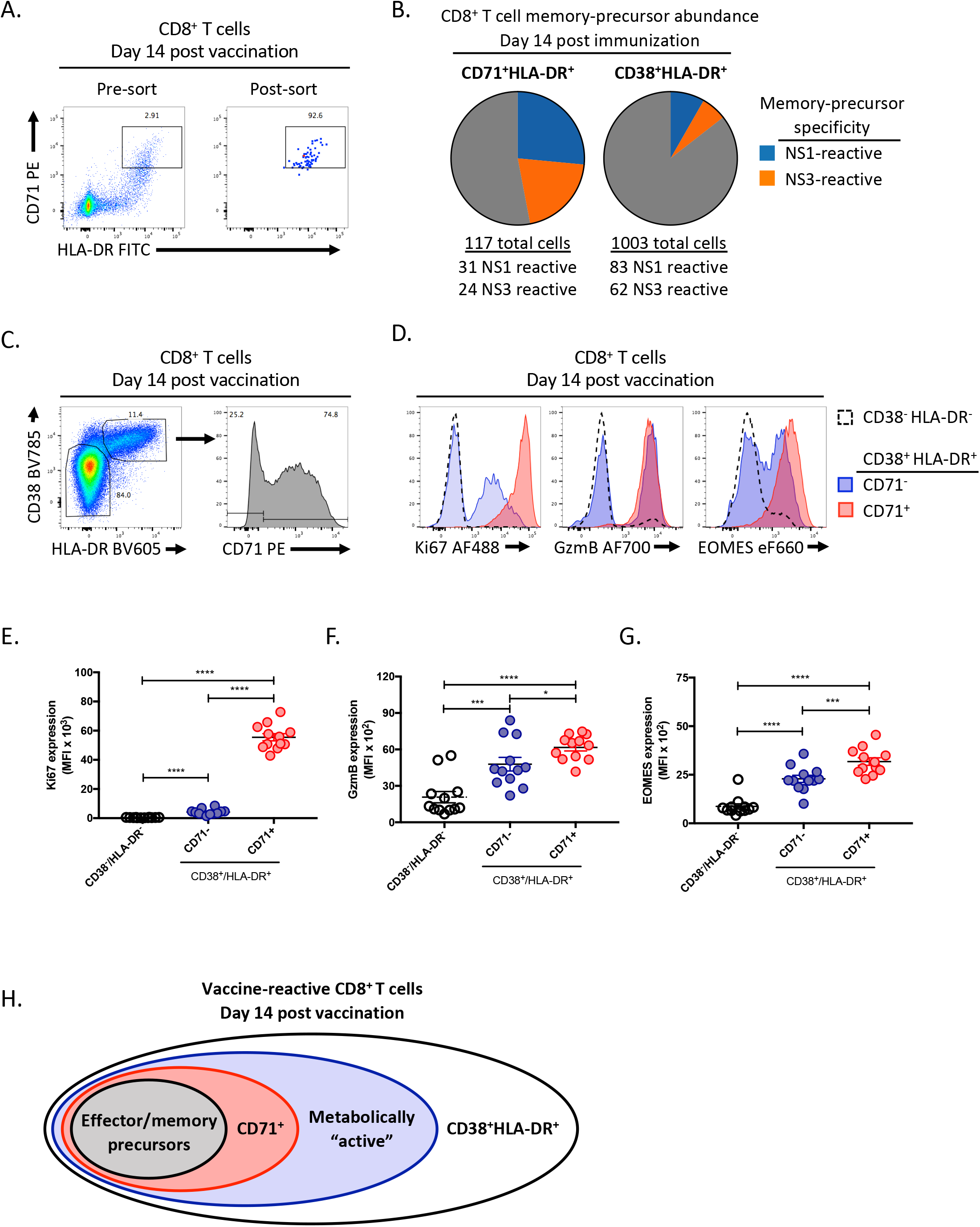
CD71 expression correlates with CD8^+^ T cell effector/memory potential in vaccine-reactive CD8^+^ T cells. CD8^+^ T cells from TAK-003-inoculated individuals were analyzed 14 days after immunization. **A)** CD71^+^HLA-DR^+^ CD8^+^ T cells were sorted from a TAK-003 inoculated individual 14 days post immunization and subjected to single-cell RNAseq analysis. **B)** The abundance of memory-precursor clonotypes was assessed at day 14 post-vaccination within the CD71^+^HLA-DR^+^ or CD38^+^HLA-DR^+^ CD8^+^ T cell compartments. Memory-precursor clonotypes were defined as TCR clones found at both day 14 within the pools of phenotypically activated CD8^+^ T cells, and at day 120 within either NS1 or NS3 reactive memory CD8^+^ T cell populations. **C)** Flow cytometric analysis of TAK-003-reactive CD8^+^ T cells 14 days post-vaccination. Cells were subdivided into CD38^-^HLA-DR^-^, and CD38^+^HLA-DR^+^ CD71^-^, and CD38^+^HLA-DR^+^CD71^+^ populations, then assessed for intracellular expression of **D)** Ki67, GzmB, and EOMES. CD71 expression on CD8^+^ T cells corresponds with significantly higher intracellular levels of **E)** Ki67, **F)** GzmB, and **G)** EOMES. Error bars show mean and SEM. N = 12 individuals. **H)** Graphical representation of the association between activation marker expression, metabolic activity, and effector/memory potential within vaccine-reactive CD8^+^ T cells. * p <0.05, *** p<0.001, **** p<0.0001 (Paired t-test)

To extend the observation that CD71 expression may better define vaccine-reactive CD8^+^ T cells with effector/memory potential, we further analyzed CD8^+^ T cells from an additional 12 individuals 14 days after immunization with TAK-003 by flow cytometry with the addition of intracellular markers of CD8^+^ T cell effector function, proliferation, and effector/memory potential. As expected, only a subset of CD38^+^HLA-DR^+^ CD8^+^ T cells express the transferrin receptor (CD71) 14 days post-TAK-003 administration (Figure 7C, **Supplemental Figure 8**). Additionally, we observed that CD71 expression within the CD38^+^HLA-DR^+^ CD8^+^T cell compartment positivity correlates with the presence of markers of cellular proliferation (Ki67) (Figure 7D, Figure 7E), cytolytic function (Granzyme B) (Figure 7D, Figure 7F), and effector/memory lineage commitment (EOMES) (Figure 7D, Figure 7G). Expression of these markers were significantly enriched in CD71^+^CD38^+^HLA-DR^+^ CD8^+^ T cells relative to CD71^-^ CD38^+^HLA-DR^+^ CD8^+^ T cells, or CD38^-^HLA-DR^-^ CD8^+^ T cells. These data demonstrate that the surface expression of CD71 is a marker of cytolytic cellular functionality and effector/memory potential, and can further aid in the rapid identification and characterization of vaccine-reactive T cells within the total “activated” CD8^+^ T cell pool elicited by vaccine exposure (Figure 7H).

## DISCUSSION

In this study, we demonstrate that the live-attenuated tetravalent DENV vaccine TAK-003 is capable of eliciting a potent and durable cellular immune response following inoculation. CD8^+^ T cell activation in response to TAK-003 administration was observed to peak 28 days post-vaccination, while maximal CD4^+^ T cell expansion occurred on day 14. DENV-specific cellular immunity persisted for at least 120 days following immunization as assessed by IFN-γ ELISPOT. The antigenic specificity of the cellular memory immune response elicited by TAK-003 spanned the entire DENV proteome and exhibited significant cross-reactivity against all four DENV serotypes. Analysis of the clonotypic and functional diversity of TAK-003-stimulated CD8^+^ T cells 14 days after vaccination utilizing scRNAseq revealed a significant amount of transcriptional heterogeneity within a phenotypically homogenous population. Isolation and scRNAseq-based analysis demonstrated that the dominant TCR clones within the NS1- and NS3-reactive memory CD8^+^ T cell populations assessed 120 days post-vaccination can also be observed within the activated CD38^+^HLA-DR^+^CD8^+^ T cell compartment 14 days post-vaccination. scRNAseq-based analysis of these “memory precursor” cells present at day 14 post-vaccination revealed a unique transcriptional signature, dominated by the gene expression pathways associated with cellular metabolism and proliferation.

Based on these observations, we were able to develop a panel of markers to assess the metabolic potential of both CD8^+^ and CD4^+^ T cells following *in vitro* or *in vivo* activation. These readily quantifiable metabolic parameters are sensitive and flexible markers of T cell activation and are easily adapted to standard flow cytometry-based immunoassays. Furthermore, we were able to demonstrate that surface expression of CD71 (transferrin receptor) marks cells with the highest functional and proliferative capacity, as assessed by Ki67, Granzyme B, and EOMES expression, providing a robust marker for identifying the CD8^+^ T cells shortly after vaccination that are most likely to contribute to a long-lived memory pool. These data not only provide insight into molecular mechanisms responsible for regulating memory T cell development, but also suggest possible therapeutic targets for enhancing vaccine efficacy by selectively priming the metabolism of effector/memory precursor CD8^+^ T cells during the critical para-vaccine T cell expansion phase.

The regulation of memory T cell development and homeostasis is a complex and incompletely-understood process, involving the integration of a constellation of immunological cues such as antigen density [41], TCR/peptide/MHC affinity [42], duration of antigen exposure [43], and cytokine availability [44–47]. However, it is becoming increasingly clear that the development of a stable memory T cell population – as well as T cell effector function and homeostasis in general – is critically dependent on the availability of a handful of key metabolites and the expression of a corresponding metabolic cellular program [48–50]. In particular, the availability of glucose [51, 52], long-chain fatty acids [53, 54], amino acids [55–57], and micronutrients such as iron [58, 59] can profoundly impact CD8^+^ T cell effector and/or memory potential, in consort with other traditional extrinsic immunological cues. Depriving T cells of any of these key metabolites either through pharmacological or genetic means has significant implications for T cell development, effector function, and long-term persistence.

Directly manipulating the metabolism of T cells *in vitro* or *in vivo* to influence effector function or persistence has primarily focused on restricting or enhancing access to the metabolite glucose. Due to the unique metabolic requirements of nascently activated T cells, which overwhelmingly eschew conventional mitochondrial oxidative phosphorylation in favor of oxidative glycolysis [48–50], glucose metabolism is a convenient therapeutic target. The proliferation and terminal effector function of both CD4^+^ and CD8^+^ T cells can be significantly enhanced by increasing glucose bioavailability [52, 60], whereas restricting glucose metabolism can facilitate the development of long-lived and/or suppressive T cell lineages [51, 61]. However, the utility of manipulating systemic glucose metabolism for therapeutic immunoregulatory effect *in vivo* is questionable. Systemic glucose metabolism is tightly regulated, and even modest perturbations in systemic glucose availability can have profound negative consequences for the well-being of patients. However, the selective upregulation of CD71 (transferrin receptor) on vaccine-reactive CD8^+^ T cells suggests that manipulating iron availability following vaccination may selectively enhance the expansion of the most functional vaccine-reactive CD8^+^ T cells. Indeed, loss-of-function mutations of the transferrin receptor have been observed and are associated with severe defects in adaptive immune function, underscoring the importance of transferrin-mediated iron uptake in maintaining functional immunity [38].

The findings presented in this study suggest that in the setting of vaccination the preferential survival of T cells undergoing clonal expansion *in vivo* is dependent on metabolite availability and the initiation of a transcriptional program permissive to nutrient uptake. Cell surface expression of the transporters of these metabolites, such as the transferrin receptor (CD71), during the acute phase proximal to vaccination, proved to be an effective marker of memory precursor CD8^+^ T cells. Cumulatively, these data highlight the utility of high content single-cell transcriptomic analysis coupled with more traditional cellular immune monitoring in assessing vaccine-elicited T cell immunity. The ability to accurately and longitudinally track T cell clones from acute infection to stable memory provides a unique opportunity to identify correlates of T cell-mediated immunity with single-cell resolution. Future work is required to verify the distinctiveness of the metabolic and transcriptional programming that determines log-term T cell memory and whether these markers define the protective capacity of T cell immunity.

## METHODS

### Cells/samples

The samples used in this study were collected during a Phase 1 trial in US adults of a tetravalent, live-attenuated dengue virus vaccine candidate, TAK-003 (NCT01728792; WRAIR #1987), as well as from healthy US adult volunteers (WRAIR #1868). TAK-003 samples were provided by Takeda Vaccines. Whole blood was collected in Cell Preparation Tubes (BD Vacutainer) for isolation of PBMC. Cells were cryopreserved at approximately 10^7^ per mL and stored in vapor-phase liquid nitrogen until use. Vaccine administration and PBMC collection were performed after written informed consent. The studies were approved by the institutional review boards at the State University of New York Upstate Medical University and the Human Subjects Research Review Board for the Commanding General of the U.S. Army Medical Research and Material Command.

### T cell ELISpot assay

Cryopreserved PBMC were thawed and placed in RPMI 1640 medium supplemented with 10% heat-inactivated normal human serum (100-318, Gemini Bio-Products), L-glutamine, penicillin, and streptomycin. After an overnight rest at 37°C, the PBMC were washed, resuspended in serum-free medium (SFM; X-VIVO 15, Lonza), and 1-2×10^5^ cells were plated per well of a 96-well Millipore MAIPSWU plate coated with anti-IFNγ antibody according to the manufacturer’s instructions (3420-2HW-Plus, Mabtech Inc.). Peptide pools were added to the cells at a final concentration of 1μg/mL/peptide prior to incubation at 37°C overnight. Controls included SFM plus 0.5% DMSO (negative) and anti-CD3 (positive). The ELISpot plates were developed using TMB substrate and read using a CTL-ImmunoSpot^®^ S6 Ultimate-V Analyzer (Cellular Technology Limited).

### Peptides

Overlapping peptide pools corresponding to the full-length envelope (E), non-structural 1 (NS1), NS2, NS3, NS4, and NS4 proteins for DENV-1-4 were obtained through the NIH Biodefense and Emerging Infections Research Resources Repository, NIAID, NIH (**Supplemental Table 2**). Additional overlapping peptide pools covering the capsid (C) and precursor membrane (prM) proteins of DENV-1-pp65 protein from hCMV, nucleocapsid protein from influenza H3N2, hexon protein from adenovirus serotype 3, and Large Envelope protein from HBV were purchased from JPT Peptide Technologies (**Supplemental Table 2**). Peptide pool stocks were reconstituted in DMSO at a concentration of 200μg/mL/peptide and stored at −80°C.

### Flow cytometry

Surface staining for flow cytometry analysis was performed in PBS supplemented with 2% FBS at room temperature. Aqua Live/Dead (ThermoFisher, L34957) was used to exclude dead cells in all experiments. Intracellular protein staining was performed using the Foxp3 Fixation/Permeabilization kit (ThermoFisher, 00-5523-00) according to the manufacturer’s recommendation. Antibodies used for flow cytometry analysis are listed in **Supplemental Figure 3**. Flow cytometry analysis was performed on a custom-order BD LSRFortessa instrument and analyzed using FlowJo v10.2 software (Treestar). Cell sorting was performed on a BD FACSAria Fusion instrument.

### *in vitro* T cell stimulation

Polyclonal T cell activation was performed by stimulating healthy donor PBMCs at a concentration of 5×10^6^ cells/mL with 0.1 μg/mL αCD3 (Clone OKT3, Biolegend 317315) and 1 μg/mL αCD28 (Clone CD28.2, BD 555725) in complete cell culture media consisting of RPMI 1640 supplemented with 10% heat-inactivated fetal bovine serum (Sigma), 2 mM L-glutamine (Lonza/BioWhittaker), and 100 U/ml penicillin/streptomycin (Lonza/BioWhittaker). For antigen-specific T cell simulation, healthy donor PBMCs were resuspended at a concentration of 5×10^6^ cells/mL in complete cell culture media and stimulated with the indicated peptide pool at a final concentration of 1 μg/mL.

### Identification and isolation of DENV-reactive memory CD8^+^ T cells

NS1- and NS3-reactive memory CD8^+^ T cells were identified and isolated from PBMC samples obtained 120 days post-vaccination. Cryopreserved PBMC samples were thawed and resuspended in complete cell culture media at a concentration of 5×10^6^ cell/mL and stimulated with 1μg/mL of NS1- or NS3-derived peptide pools (**Supplemental Table 2**) for 18 hours at 37° C. NS1- or NS3-reactive CD8^+^ T cells were identified by expression of the activation markers CD25 and CD69, and isolated by flow sorting.

### Metabolite uptake assay

Cryopreserved PBMC samples were thawed and resuspended in complete cell culture media at a concentration of 5×10^6^ cell/mL and rested for 30 min at 37° C prior to metabolic analysis. For assessment of glucose uptake, cells were subsequently washed 2X with glucose-free RPMI (ThermoFisher, N13195), resuspended at a concentration of 5×10^6^ cell/mL in glucose-free RPMI, and rested for an additional 10 minutes at 37° C. 2-NBDG (ThermoFisher, D3821) was added to a final concentration of 100μM, and cells were incubated for 30 minutes at 37° C. Cells were washed 2X with PBS + 5% FBS, then surface stained for flow cytometry analysis as described above. For assessment of fatty-acid uptake, BODIPY FL-C_16_ was added to cells in complete cell culture media at a final concentration of 1μM. Cells were incubated for 30 minutes at 37° C, then washed 2X with PBS + 5% FBS and surfaced stained for flow cytometry analysis as described above.

### Single-cell RNA sequencing library generation

Flow-sorted CD8^+^ T cell suspensions at a density of 50-500 cells/μL in PBS plus 0.5% FBS were prepared for single-cell RNA sequencing using the Chromium Single Cell 5’ Reagent version 2 kit and Chromium Single Cell Controller (10x Genomics, CA) as previously described with some modification [42]. In short, 500 – 2,000 cells per reaction were loaded for gel bead-in-emulsion (GEM) generation and barcoding. Reverse transcription, RT-cleanup, and cDNA amplification were performed to isolate and amplify cDNA for downstream 5’ gene or enriched V(D)J library construction according to the manufacture’s protocol. Libraries were constructed using the Chromium Single Cell 5’ reagent kit, V(D)J Human T Cell Enrichment Kit, 3’/5’ Library Construction Kit, and i7 Multiplex Kit (10x Genomics, CA) according to the manufacture’s protocol.

### Sequencing

scRNAseq 5’ gene expression libraries were sequenced on an Illumina NextSeq platform with a 500/550 High Output Kit v2 (150 cycles) to a read depth of ~30,000 reads/cell. Sequencing parameters were set for Read1 (26 cycles), Index1 (8 cycles), and Read2 (98 cycles). scRNAseq TCR V(D)J enriched library sequencing was performed on an Illumina MiSeq platform with a v3 Reagent Kit (600 cycles) to a read depth of ~10,000 reads/cell. Sequencing cycles were set at 150 for Read1, 8 for Index1), and 150 for Read2. Prior to sequencing, library quality and concentration were assessed using an Agilent 4200 TapeStation with High Sensitivity D5000 ScreenTape Assay and Qubit Fluorometer (Thermo Fisher Scientific) with dsDNA BR assay kit according to the manufacturer’s recommendation.

### 10x Genomics 5’ gene-expression data analysis

5’ gene expression analysis from sorted CD8^+^ T cells was performed using the 10x Genomics Cell Ranger pipeline as previously described [42]. In short, sample demultiplexing and analysis was performed using the Cell Ranger software package (10x Genomics, CA, v2.1.0) according to the manufacturer’s recommendations, with the default settings, and *mkfastq/count* commands, respectively. Transcript alignment was performed against a human reference library generated using the Cell Ranger *mkref* command and the Ensembl GRCh38 v87 top-level genome FASTA and the corresponding Ensembl v87 gene GTF. Data visualization and differential gene expression analysis were performed using the Loupe Cell Browser (10x Genomics, CA, v2.0.0). t-SNE plot visualization of gene expression data was based on the cellular coordinates calculated by the Cell Ranger *count* command.

### TCR sequence analysis

Sorted CD8^+^ T cell TCR clonotype identification, alignment, and annotation was performed using the 10x Genomics Cell Ranger pipeline. Sample demultiplexing and clonotype alignment was performed using the Cell Ranger software package (10x Genomics, CA, v2.1.0) according to the manufacturer’s recommendations, with the default settings, and *mkfastq/vdj* commands, respectively. TCR clonotype alignment was performed against a filtered human V(D)J reference library generated using the Cell Ranger *mkvdjref* command and the Ensembl GRCh38 v87 top-level genome FASTA and the corresponding Ensembl v87 gene GTF. TCR clonotype visualization, diversity assessment, and analysis were performed using the Loupe VDJ Browser (10x Genomics, CA, v2.0.0). TCR gene segment usage was assessed and visualized using VDJTools [62].

### Statistical analysis

All statistical analysis was performed using GraphPad Prism 6 Software (GraphPad Software, La Jolla, CA). A p-value <0.05 was considered significant.

## Acknowledgments

This work was supported by the Military Infectious Disease Research Program (MIDRP) and the Congressionally Directed Medical Research Program (CDMRP). This research was performed while Dr. Waickman held an NRC Research Associate award at the Walter Reed Army Institute of Research.

## Disclaimer

The opinions or assertions contained herein are the private views of the authors and are not to be construed as reflecting the official views of the US Army or the US Department of Defense. Material has been reviewed by the Walter Reed Army Institute of Research. There is no objection to its presentation and/or publication. The investigators have adhered to the policies for protection of human subjects as prescribed in AR 70–25.

