## Supplemental Figures for "Dissecting the heterogeneity of DENV vaccine-elicited cellular immunity using single-cell RNA sequencing and cellular metabolic profiling"

### Supplemental Figure 1

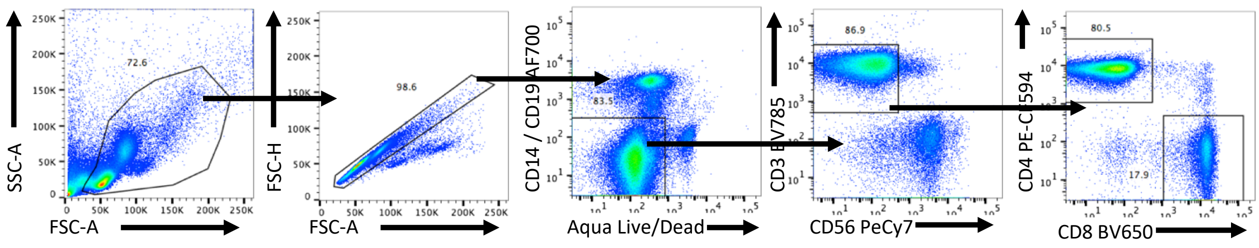


**Supplemental figure 1: Gating strategy for Figure 1A and Figure 1C.**

#### Supplemental Figure 2

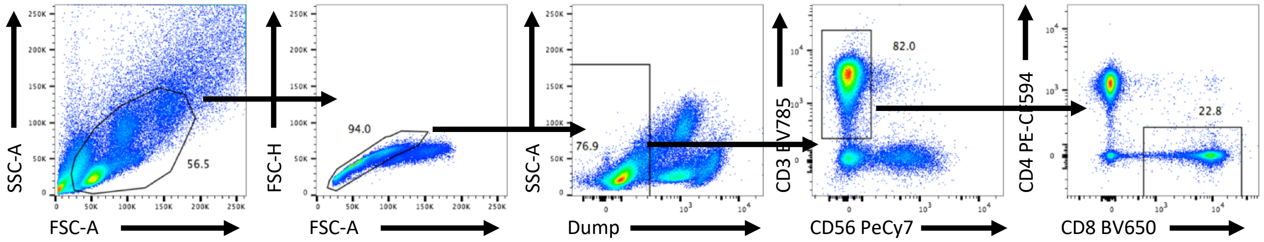


**Supplemental figure 2: Gating strategy for Figure 2B and Figure 2C.** Dump channel contains viability marker (Aqua Live/Dead), CD14 BV510, and CD19 BV510.

**Supplemental figure 3**

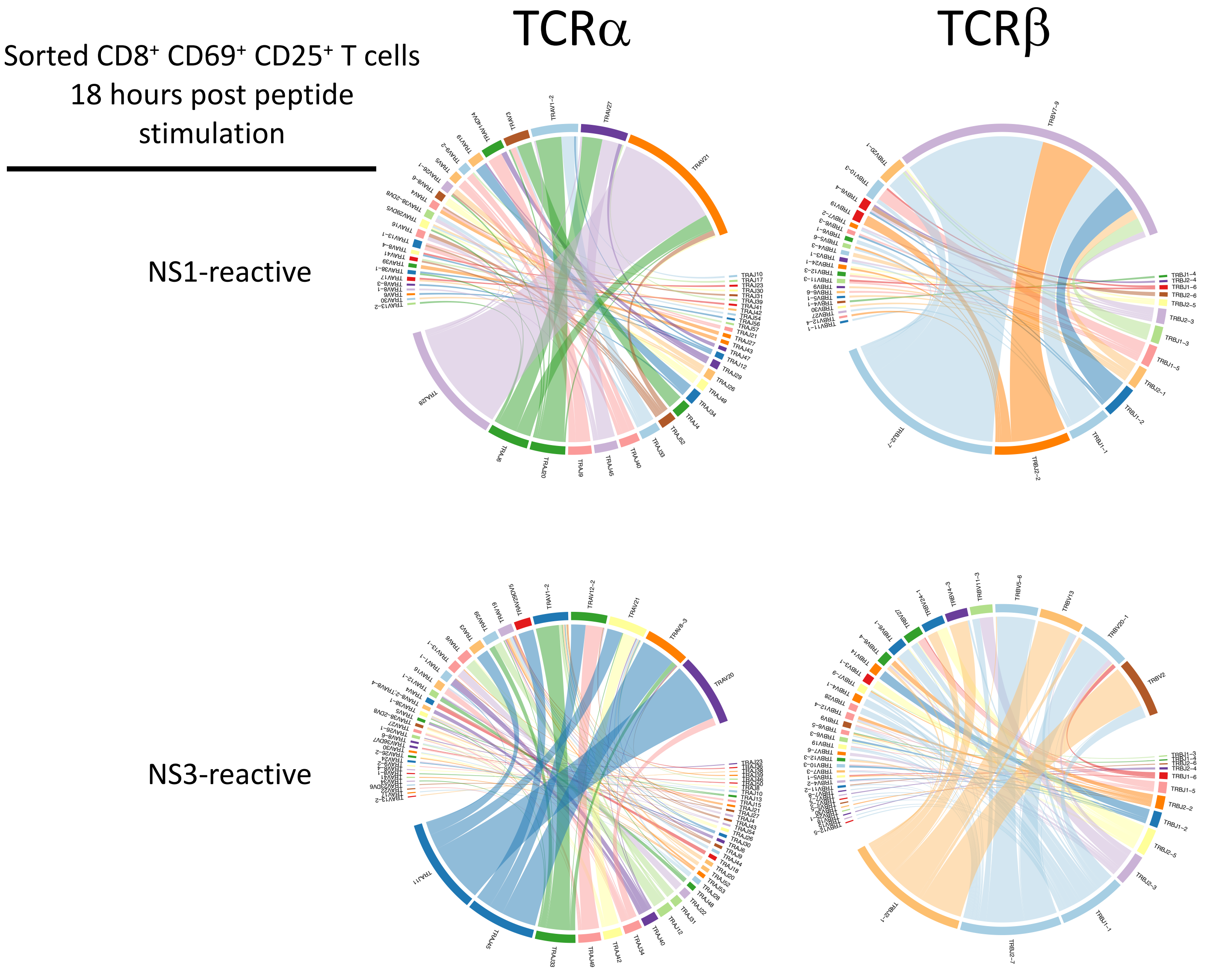


**Supplemental figure 3: TCRα and TCRβ V/J segment usage and pairing from sorted NS1- and NS3-reactive CD8+ memory T cells 120 days after TAK-003 administration.**

### Supplemental figure 4

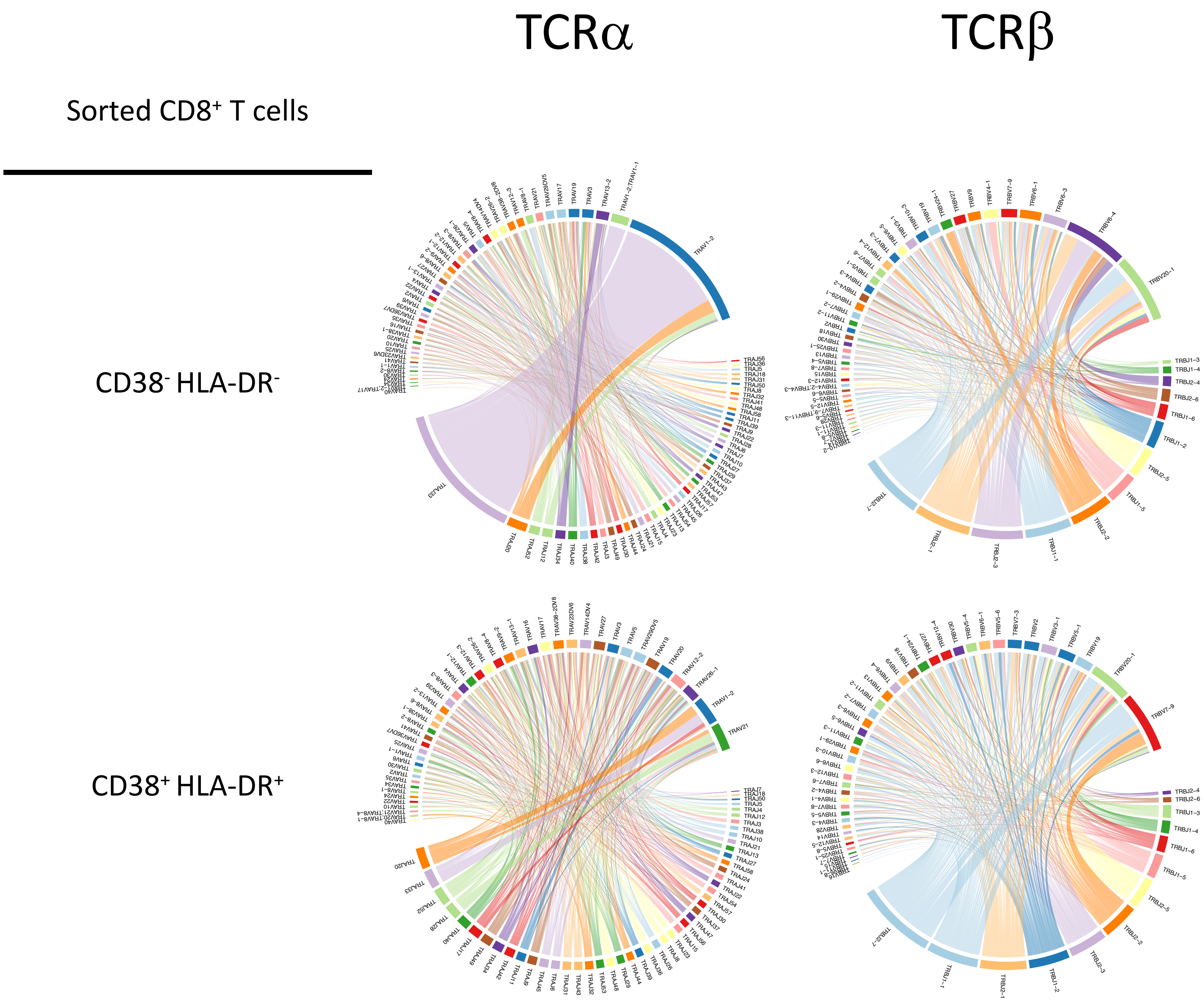


**Supplemental figure 4: TCRα and TCRβ V/J segment usage and pairing from sorted CD38⁻HLA-DR⁻ CD8⁺ T cells and CD38⁺HLA-DR⁺ CD8⁺ T cells 14 days post TAK-003 administration.**

### Supplemental Figure 5

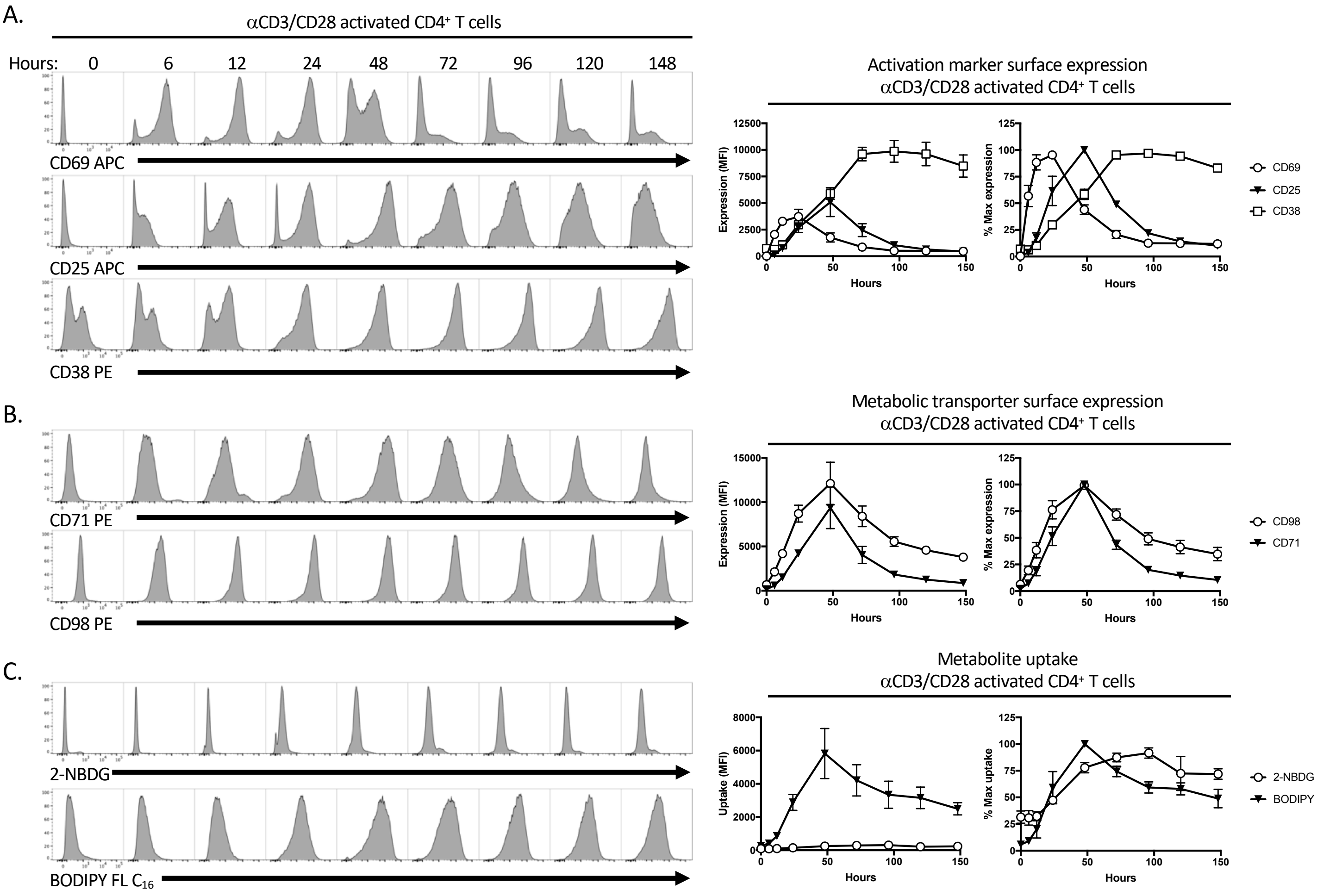


**Supplemental figure 5:** Metabolic marker upregulation on *in vitro* stimulated CD4 T cells. PBMCs from healthy donors were stimulated with 0.1mg/ml aCD3 and 1mg/ml aCD28 and expression of **A)** CD69 and CD25 **B)** CD71 and CD98, or **C)** uptake of 2-NBDG and BODIPY FL-$C_{16}$ at the indicated timepoints. Results are representative of two independent experiments, with a total of 4 individual donors.

### Supplemental figure 6

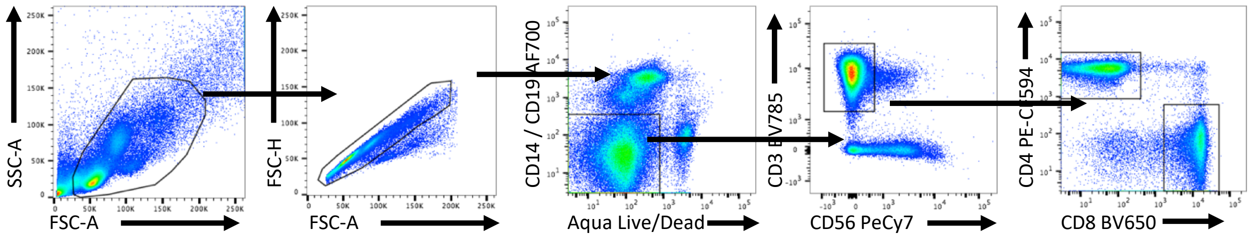


**Supplemental figure 6:** Gating scheme scheme for Figure 6 and Supplemental Figure7

### Supplemental figure 7

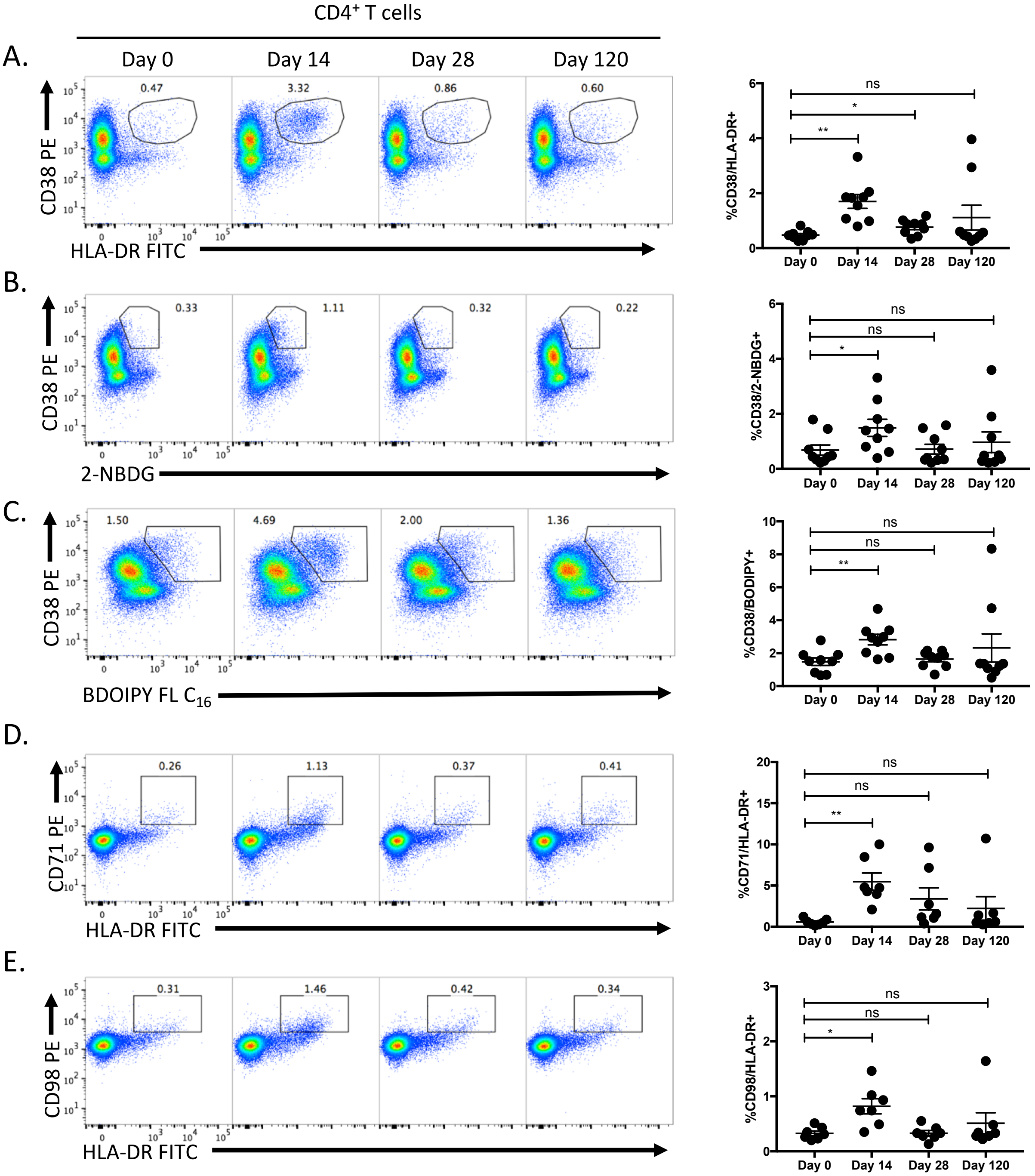


**Supplemental figure 7: Vaccine-reactive CD4+ T cells are identifiable by changes in metabolite transporter expression and metabolite utilization**. CD4 T cells from TAK-003 recipients were analyzed by flow cytometry at days 0, 14, 28 and 120 post vaccination. Vaccine-reactive CD4+ T cells were quantified based on expression of **A)** CD38/HLA-DR, **B)** CD38/2-NBDG, **C)** CD38/BODIPY FL-C16, **D)** CD71/HLA-DR, or **E)** CD98/HLA-DR. Error bars show mean and SEM. N = 10 individuals. * $p < 0.05$, ** $p < 0.01$ (paired t test)

### Supplemental Figure 8

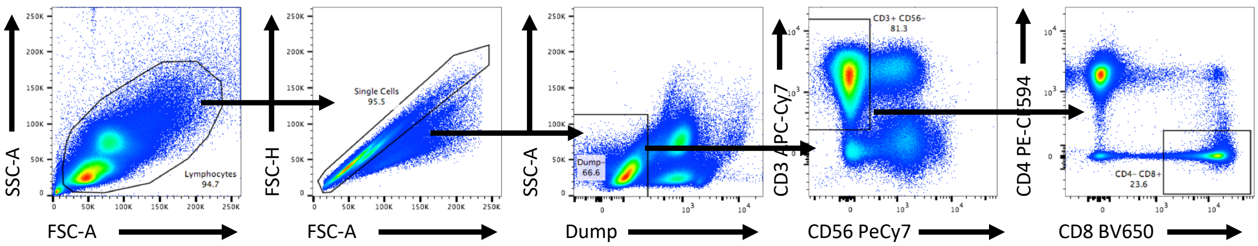


**Supplemental figure 8: Gating strategy for Figure 7.** Dump channel contains viability marker (Aqua Live/Dead), CD14 BV510, and CD19 BV510.
