## Supplemental Tables for "Dissecting the heterogeneity of DENV vaccine-elicited cellular immunity using single-cell RNA sequencing and cellular metabolic profiling"

**Supplemental Table 1.** Dominant TCR clones from CD71$^+$HLA-DR$^+$ CD8$^+$ T cells isolated 14 days post DENVax administration

| | Day 14, CD71$^+$HLA-DR$^+$ | | | | | | |
|---|---|---|---|---|---|---|---|
| | TCRα | | | TCRβ | | | |
| **Frequency** | **CDR3aa** | **V** | **J** | **CDR3aa** | **V** | **D** | **J** |
| 0.1 | CAPLGGAGSYQLTF | TRAV21 | TRAJ28 | CASSPRQGNTGELFF | TRBV7-9 | TRBD1 | TRBJ2-2 |
| 0.075 | CAGRGAGSYQLTF | TRAV21 | TRAJ28 | CASSLLSYEQYF | TRBV7-9 | | TRBJ2-7 |
| 0.0625 | CAVRFPRDYKLSF | TRAV21 | TRAJ20 | CASSPTGTGYYEQYF | TRBV7-9 | TRBD1 | TRBJ2-7 |
| 0.05 | CAVRGRGDYKLSF | TRAV1-2 | TRAJ20 | CASSSAGTLNTGELFF | TRBV7-9 | TRBD1 | TRBJ2-2 |
| 0.0375 | CAGAWKNTGFQKLVF | TRAV27 | TRAJ8 | CASSEWEGNYGYTF | TRBV6-1 | TRBD2 | TRBJ1-2 |
| 0.0375 | CALSEAQYNFNKFYF | TRAV19 | TRAJ21 | CASSIPTSGTLGDTQYF | TRBV5-6 | TRBD2 | TRBJ2-3 |
| 0.0375 | CIVRSLINYGQNFVF | TRAV26- | TRAJ26 | CASSSVSYEQYF | TRBV7-9 | | TRBJ2-7 |
| 0.0375 | CAVQAGGYSTLTF | TRAV20 | TRAJ11 | CASAEADNEQFF | TRBV13 | TRBD2 | TRBJ2-1 |
| 0.025 | CAVNEAGGFKTIF | TRAV12- | TRAJ9 | CASSLEVFDEQFF | TRBV5-6 | | TRBJ2-1 |
| 0.025 | CAESGDSNYQLIW | TRAV5 | TRAJ33 | CAWSVGGTGELFF | TRBV30 | TRBD1 | TRBJ2-2 |

**Supplemental table 2.** Peptide pools utilized in this study

| DENV type | Strain | Protein region | Size (#aa) | Overlap (#aa) | Source | Cat No. |
|---|---|---|---|---|---|---|
| DENV-1 | Nauru/West Pac/1974 | C/prM | 16 | 11 | JPT Peptide Technologies | custom |
| | Nauru/West Pac/1974 | E | 13-18 | 11-12 | BEI Resources | NR-9241/ NR-4551 |
| | Singapore/S275/1990 | NS1 | 13-17 | 11-12 | BEI Resources | NR-2751 |
| | Singapore/S275/1990 | NS3 | 14-17 | 11-12 | BEI Resources | NR-2752 |
| | Singapore/S275/1990 | NS5 | 12-17 | 11-12 | BEI Resources | NR-4203 |
| DENV-2 | S16803 | C/prM | 16 | 11 | JPT Peptide Technologies | custom |
| | New Guinea C | E | 15-20 | 10-11 | BEI Resources | NR-507 |
| | New Guinea C | NS1 | 15-19 | 10-11 | BEI Resources | NR-508 |
| | New Guinea C | NS2a | 15-17 | 11 | BEI Resources | NR-2747 |
| | New Guinea C | NS2b | 13-17 | 11-14 | BEI Resources | NR-2748 |
| | New Guinea C | NS3 | 13-19 | 10 | BEI Resources | NR-509 |
| | New Guinea C | NS4a | 14-17 | 11-12 | BEI Resources | NR-2749 |
| | New Guinea C | NS4b | 12-17 | 11 | BEI Resources | NR-2750 |
| | New Guinea C | NS5 | 15-17 | 11-13 | BEI Resources | NR-2746 |
| DENV-3 | CH53489 | C/prM | 16 | 11 | JPT Peptide Technologies | custom |
| | Philippines/H87/1956 | E | 12-20 | 10-11 | BEI Resources | NR-9228/ NR-511 |
| | Philippines/H87/1956 | NS1 | 13-17 | 11-12 | BEI Resources | NR-2753 |
| | Philippines/H87/1956 | NS3 | 14-17 | 11-12 | BEI Resources | NR-2754 |
| | Philippines/H87/1956 | NS5 | 13-17 | 11-13 | BEI Resources | NR-4204 |
| DENV-4 | Singapore/8976/1995 | C/prM | 16 | 11 | JPT Peptide Technologies | custom |
| | Singapore/8976/1995 | E | 12-20 | 10-11 | BEI Resources | NR-9229/ NR-512 |
| | Singapore/8976/1995 | NS1 | 13-17 | 11-12 | BEI Resources | NR-2755 |
| | Singapore/8976/1995 | NS3 | 15-17 | 11-12 | BEI Resources | NR-2756 |
| | Singapore/8976/1995 | NS5 | 13-17 | 11-14 | BEI Resources | NR-4205 |

**Supplemental Table 3.** Antibodies utilized for flow cytometry analysis in this study

| Antibody | Manufacture | Clone | Cat# |
|---|---|---|---|
| CD14 AF700 | BD | M5E2 | 557923 |
| CD14 BV510 | BD | MOP9 | 563079 |
| CD19 AF700 | BD | HIB19 | 557921 |
| CD19 BV510 | BD | SJ25C1 | 562947 |
| CD56 PE-Cy7 | BD | B159 | 557747 |
| CD98 PE | BD | UM7F8 | 556077 |
| Granzyme B AF700 | BD | GB11 | 560213 |
| HLA-DR FITC | BD | G46-6 | 555811 |
| Ki67 AF488 | BD | B56 | 562900 |
| CD3 BV785 | Biolegend | OKT3 | 317330 |
| CD38 BV421 | Biolegend | HIT2 | 303526 |
| CD38 PE | Biolegend | HB-7 | 356604 |
| CD4 PE-Dzz594 | Biolegend | RPA-T4 | 300548 |
| CD69 APC | Biolegend | FN50 | 310910 |
| CD69 APC-Cy7 | Biolegend | FN50 | 310914 |
| CD71 PE | Biolegend | CY1G4 | 334106 |
| CD8 BV650 | Biolegend | RPA-T8 | 301042 |
| HLA-DR BV605 | Biolegend | L243 | 307640 |
| CD25 APC | Invitrogen | BC96 | 17-0259-42 |
| EOMES eF660 | Invitrogen | WD192B | 50-4877-42 |
